# The apicoplast is important for the viability and persistence of *Toxoplasma gondii* bradyzoites

**DOI:** 10.1101/2023.01.13.523885

**Authors:** Syrian G. Sanchez, Emilie Bassot, Aude Cerutti, Hoa Mai Nguyen, Amel Aïda, Nicolas Blanchard, Sébastien Besteiro

**Author notes:** Corresponding author: Sébastien Besteiro. **Author Contributions**: N. Blanchard and S. Besteiro designed research; S. Sanchez, E. Bassot, A. Cerutti, H.M. Nguyen, A. Aïda S. Besteiro, performed research; S. Sanchez, E. Bassot, N. Blanchard and S. Besteiro analyzed the data; S. Sanchez, N. Blanchard and S. Besteiro wrote the paper; N. Blanchard and S. Besteiro acquired funding.

## Abstract

*Toxoplasma gondii* is responsible for toxoplasmosis, a disease that can be serious when contracted during pregnancy, but can also be a threat for immunocompromised individuals. Acute infection is associated with the tachyzoite form that spreads rapidly within the host. However, under stress conditions, some parasites can differentiate into cyst-forming bradyzoites, residing mainly in the central nervous system, retina and muscle. Because this latent form of the parasite is resistant to all currently available treatments, and is central to persistence and transmission of the parasite, new specific therapeutic strategies targeting this developmental stage need to be discovered.

*T. gondii* contains a plastid of endosymbiotic origin called the apicoplast, which is an appealing drug target because it is essential for tachyzoite viability and contains several key metabolic pathways that are largely absent from the mammalian host. Its function in bradyzoites, however, is unknown. Our objective was thus to study the contribution of the apicoplast to the viability and persistence of bradyzoites during chronic toxoplasmosis.

We have used complementary strategies based on stage-specific promoters to generate conditional bradyzoite mutants of essential apicoplast genes. Our results show that specifically targeting the apicoplast in both *in vitro* or *in vivo*-differentiated bradyzoites leads to a loss of long-term bradyzoite viability, highlighting the importance of this organelle for this developmental stage. This validates the apicoplast as a potential area to look for new therapeutic targets in bradyzoites, with the aim to interfere with this currently incurable parasite stage.

**Significance Statement:** In its intermediate hosts, the parasite *Toxoplasma gondii* can persist as a cyst-contained developmental form that might reactivate and cause severe pathologies. Importantly, this form is resistant to current anti-parasitic drugs. *T. gondii* harbors a plastid of endosymbiotic origin called the apicoplast, containing important and potentially druggable metabolic pathways, but whose contribution to the fitness and viability of persistent parasites has never been assessed. We have generated conditional mutants specifically affected for the homeostasis of the apicoplast in cyst-contained parasites and showed that this organelle is important for persistence of these particular developmental forms. Our work thus validates the apicoplast as a relevant drug target in the context of chronic *T. gondii* infection.

## Introduction

*Toxoplasma gondii* is an intracellular parasitic protist responsible for toxoplasmosis, a ubiquitous disease affecting humans, which can lead to severe forms during pregnancy or in immunosuppressed individuals (1). Acute toxoplasmosis is associated with the tachyzoite form of these parasites, which spreads rapidly within the host and has the ability to cross physiological barriers, but is generally controlled by the immune system in immunocompetent hosts (2). The genus *Toxoplasma* contains only one species, but there are many different strains which are grouped according to their virulence and pathogenicity in mouse models (3). The pathogenic outcome is closely linked to the ability of the parasite to convert from fast-replicating tachyzoites to persisting encysted stages that can be found predominantly in neurons and muscle tissues and are called bradyzoites (4). These developmental forms of the parasite, which are the hallmark of chronic infection, are supposedly less active metabolically and slower to replicate than tachyzoites (5). However, they play a crucial role in *T. gondii* transmission, as well as in immune escape and survival of the parasites under stressful conditions (6).

The process of stage conversion is a continuum lasting for several days that leads to marked differences between tachyzoites and bradyzoites. For instance the membrane of the parasitophorous vacuole (PV, the compartment in which the parasite replicates) transforms into a heavily-glycosylated cyst wall (7, 8). Also, several aspects of parasite metabolism change drastically, as illustrated by the accumulation of amylopectin granules in bradyzoites (9) and the expression of different stage-specific isoforms of metabolic enzymes (10–14). Thus unsurprisingly, stage conversion is accompanied by a considerable change in protein expression driven by stage-specific promoters. Transcriptomic analyses of *in vivo*-generated cysts have for instance revealed that the expression levels of hundreds of genes are potentially changing between bradyzoites and tachyzoites (15, 16). The specific metabolic requirements of bradyzoites are so far poorly characterized, largely due to the technical challenges associated with studying this developmental stage: although *in vitro* differentiation can be achieved by applying stresses to a cystogenic *T. gondii* strain or by infecting specific host cell types, *in vitro*-derived bradyzoites are difficult to keep in long-term culture and they usually do not reach the stage of complete maturity that is observed in mouse brain-derived bradyzoites (17–20).

As intracellular developmental stages, both PV-contained tachyzoites and cyst-enclosed bradyzoites are tightly interconnected metabolically with their host cells to ensure optimal proliferation or persistence (21). While *T. gondii* relies on specific host cell resources for survival, it is also able to synthesize *de novo* a number of important metabolites. One key parasite organelle for metabolite production is the apicoplast. This plastid of secondary endosymbiotic origin has lost its photosynthetic capacity (22), but it nevertheless harbors four main metabolic pathways which have all been shown to be essential for the *in vitro* fitness of *T. gondii* tachyzoites (23, 24). These are important for the synthesis of fatty acids (FAs, via the FASII pathway), heme (together with the mitochondrion), isoprenoid precursors and iron-sulfur clusters. Because of the prokaryotic origin and the metabolic importance of most of the apicoplast-hosted pathways, this organelle is particularly attractive as a potential drug target and compounds inhibiting translation in the apicoplast, like clindamycin, are already used against acute toxoplasmosis (25).

However, to this day there is no drug available to clear the cyst form that persists during the chronic phase of infection, and this has major implications for pathogenesis. Clearly, curing patients from the infection is of paramount importance to prevent parasite reactivation. Several compounds, including atovaquone and quinolone derivatives, or even combinatorial strategies involving apicoplast targeting drugs like clindamycin and spiramycin, are able to limit cyst burden in animal models of chronic toxoplasmosis (26–32). However, most fail to completely eradicate bradyzoites, and some are even known to be potent inducers of stage conversion in vitro (33, 34). Yet, it is still unclear whether the resistance to these drugs is due to poor accessibility of brain-localized cysts or to the metabolically quiescent nature of the bradyzoites.

The metabolic contribution of the apicoplast to the viability and persistence of these developmental stages is still a major conundrum. Although Atg8, a protein that is important for maintaining apicoplast homeostasis (35) was found to be essential for bradyzoite viability, its involvement in canonical autophagy, which is also essential for the survival of this particular stage, did not allow demonstrating the essentiality of the organelle (36). To get insights into the importance of the organelle in the context of chronic *T. gondii* infection, we have thus generated conditional mutants more specifically affected in apicoplast function at the bradyzoite stage. As the apicoplast is essential to the viability of tachyzoites, which are a necessary step prior to obtaining bradyzoites, we needed to adopt a conditional strategy for targeting the organelle. Most conditional expression systems available for *T. gondii* involve the use of small molecules (37), however the bioavailability of these compounds may be difficult to regulate in the *in vivo* mouse model and most of them may not efficiently cross the blood-brain barrier to reach the tissue cysts. We thus genetically-engineered parasite strains to impact apicoplast functions specifically in bradyzoites and without the use of external compounds, by using stage-specific promoters. We generated mutants either affected for apicoplast-related metabolism or in the actual structure and division process of the organelle. Through *in vitro* and *in vivo* phenotypic analyses, we showed that in both cases the viability or development of bradyzoites was impaired. Our study thus validates the apicoplast as a potential drug target in the context of chronic toxoplasmosis.

## Results

### Specific down-regulation of the APT1 apicoplast phosphate translocator at the bradyzoite stage

To generate a parasite line impacted for apicoplast homeostasis, we first targeted the Apicoplast triosephosphate translocator 1 (APT1). APT1 is involved in the import of phosphoenol pyruvate and triose phosphates, which are important precursors for the FASII system and for isoprenoid synthesis in the organelle (Fig. 1A), and as such it is essential to the survival of *T. gondii* tachyzoites (38). Although the metabolic requirements of the bradyzoites are not completely elucidated yet, we reasoned that depleting a transporter which is needed for two of the most important apicoplast-hosted metabolic pathways would be likely to have an impact. To generate a conditional mutant specifically affected in bradyzoites, we used CRISPR/Cas9 to edit the endogenous *APT1* locus in the cystogenic Prugniaud (Pru) strain: we inserted a cassette coding for a GFP-fused version of the APT1 protein driven by a tachyzoite stage-specific *SAG1* promoter (Fig. 1B). Proper integration in the resulting cell line, named pSAG1-APT1, was verified by PCR (Fig. S1A). Using co-staining with TgAtrx1, a marker of the periphery of the apicoplast (39), we also checked that the promoter change and the addition of the GFP tag were not affecting the localization of APT1 at the organelle (Fig. S1B). However, it should be noted that the *SAG1* promoter is potentially a much stronger promoter than the native *APT1* promoter according to comparative analysis of RNA expression level (6). Accordingly, we performed semi-quantitative RT-PCR analyses on tachyzoites that confirmed at least a 3-fold increase in expression for the *APT1* mRNA upon promoter change (Fig. S1C, D).

**Figure 1.**
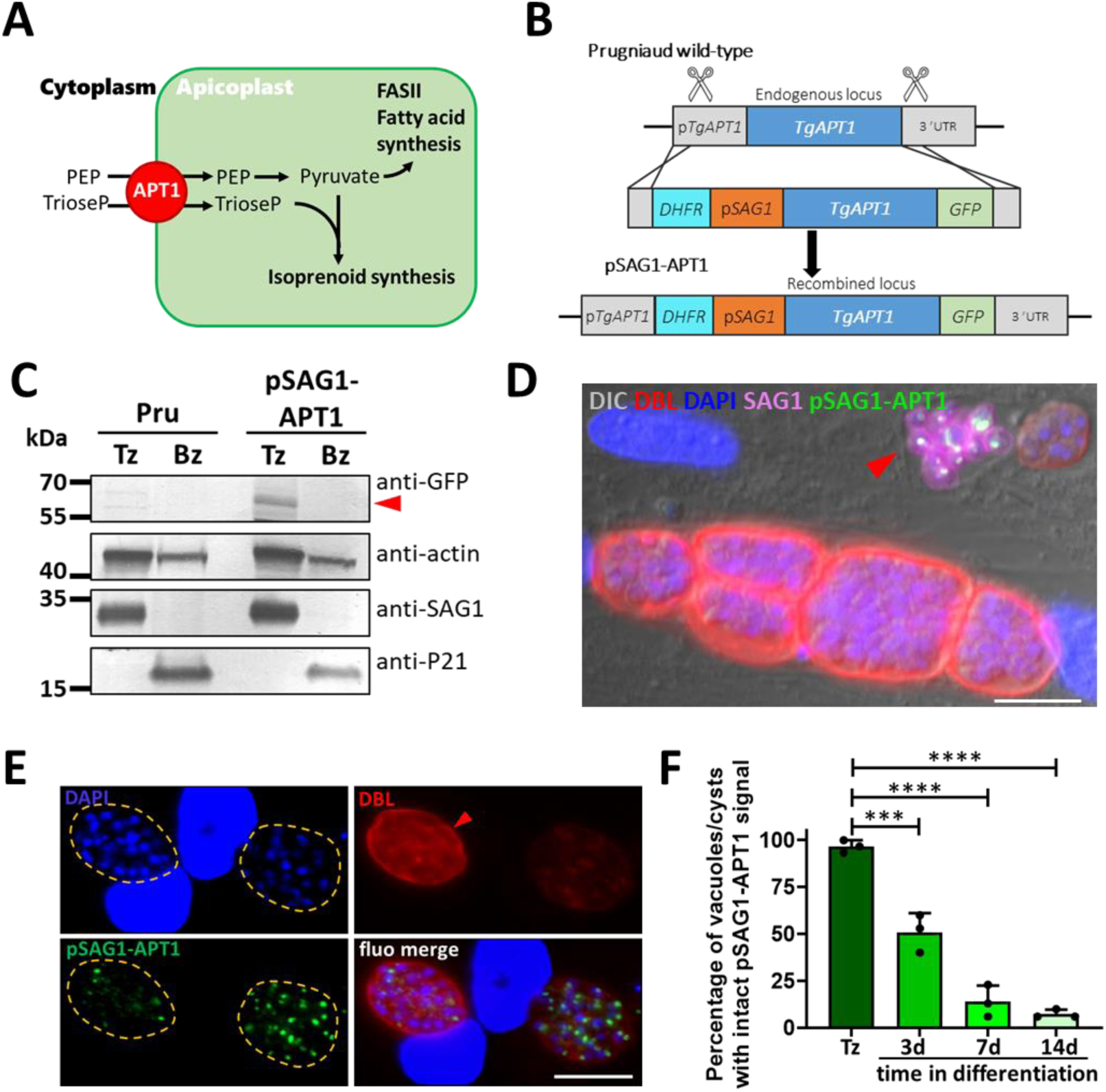
Conditional depletion of APT1 at the bradyzoite stage. A) Schematic representation of APT1 function at the apicoplast and its implication upstream of two important metabolic pathways. B) Scheme describing the strategy to edit the *APT1* locus by homologous recombination, to express a copy of the *APT1* gene under the dependence of a tachyzoite-specific *SAG1* promoter and allowing fusion of APT1 with a C-terminal GFP. C) Immunoblot analysis of protein extracts from tachyzoites (Tz) and 14 days *in vitro*-differentiated (alkaline stress) bradyzoites (Bz) with an anti-GFP antibody to detect the expression of APT1-GFP (arrowhead) under the dependence of the *SAG1* promoter. Actin was used as a loading control, SAG1 and P21 as tachyzoite and bradyzoite markers, respectively. D) Microscopic imaging of pSAG1-APT1 parasites submitted to alkaline pH stress for 7 days shows that APT1 expression (green) is coordinated with the expression of SAG1 (magenta) in non-differentiated parasites (arrowhead). Cyst walls were labelled with *D. biflorus* lectin (DBL). Scale bar=10 µm. E) Microscopic imaging of pSAG1-APT1 parasites differentiated as in D). Cyst walls were labelled with DBL or outlined with dashed lines. A more intense DBL labelling outlines a more mature cyst (arrowhead). DNA was labelled with DAPI. Scale bar=10 µm. F) The GFP-APT1 signal in vacuoles containing tachyzoites, or cysts generated by alkaline pH-induced differentiation, was quantified and shows progressive signal loss upon stage conversion. Data are mean from *n*=3 independent experiments +SD. *** *p* ≤ 0.001, **** *p* ≤ 0.0001, by one-way ANOVA followed by Tukey post-hoc test.

Conversion of tachyzoites into bradyzoites can be initiated *in vitro* by applying a variety of stresses like alkaline pH, heat shock, nutrient starvation or the use of specific drugs (4). For instance, alkaline pH stress-induced differentiation, which is performed in absence of CO_2_, is commonly used to induce conversion to bradyzoites with minimal toxicity to host cells. We thus used this stress on the pSAG1-APT1 cell line and performed immunoblot analysis that showed a decrease in expression of the *SAG1* promoter-driven APT1 protein in *in vitro*-generated bradyzoites (Fig. 1C). We performed fluorescence microscopy on differentiating parasites using a lectin from the plant *Dolichos biflorus* (DBL) that recognizes the SRS44/CST1 cyst wall glycoprotein that accumulates during differentiation (40). Strikingly, pSAG1-APT1 expression was tightly coordinated with SAG1 expression, and consequently APT1 was found to be largely absent from more mature cysts (Fig. 1D, E). Quantification showed that the proportion of cysts displaying partial or complete loss of the pSAG1-APT1 signal increased over time during pH stress-induced differentiation (Fig. 1F).

Overall, our data show that the *SAG1* promoter replacement strategy leads to an efficient down-regulation of APT1 upon *in vitro* differentiation of bradyzoites.

### Down-regulation of APT1 leads to perturbation of apicoplast homeostasis and impacts the development and viability of *in vitro*-generated bradyzoites

We next evaluated the impact of APT1 down-regulation on the overall homeostasis of the apicoplast. Depleting APT1 in tachyzoites has been shown not to have an immediate impact on the morphology of the organelle (38). However, this translocator is important for isoprenoid production, whose perturbation leads to loss of the organelle in the related malaria-causing *Plasmodium* parasites (41, 42), so we anticipated that long term depletion could also have an effect in bradyzoites. Co-staining of APT1 with TgCpn60 (a marker of the apicoplast lumen (43)) during the differentiation process showed a disappearance of both proteins in more mature cysts (Fig. 2A). Quantification showed that only about 35% of the cysts still contained an intact TgCpn60 signal after two weeks of pH stress-induced differentiation (Fig. 2B). As this could reflect a specific impact on this protein marker rather than a more general organelle defect, we also stained for another apicoplast protein, the E2 subunit of the pyruvate dehydrogenase (44) and observed a similar loss (Fig. S2). In cysts presenting defaults, both apicoplast markers were extensively lost (Figs. S2 and S3). Overall, this suggests that APT1 loss subsequently leads to a perturbation of several apicoplast proteins, if not the whole organelle.

**Figure 2.**
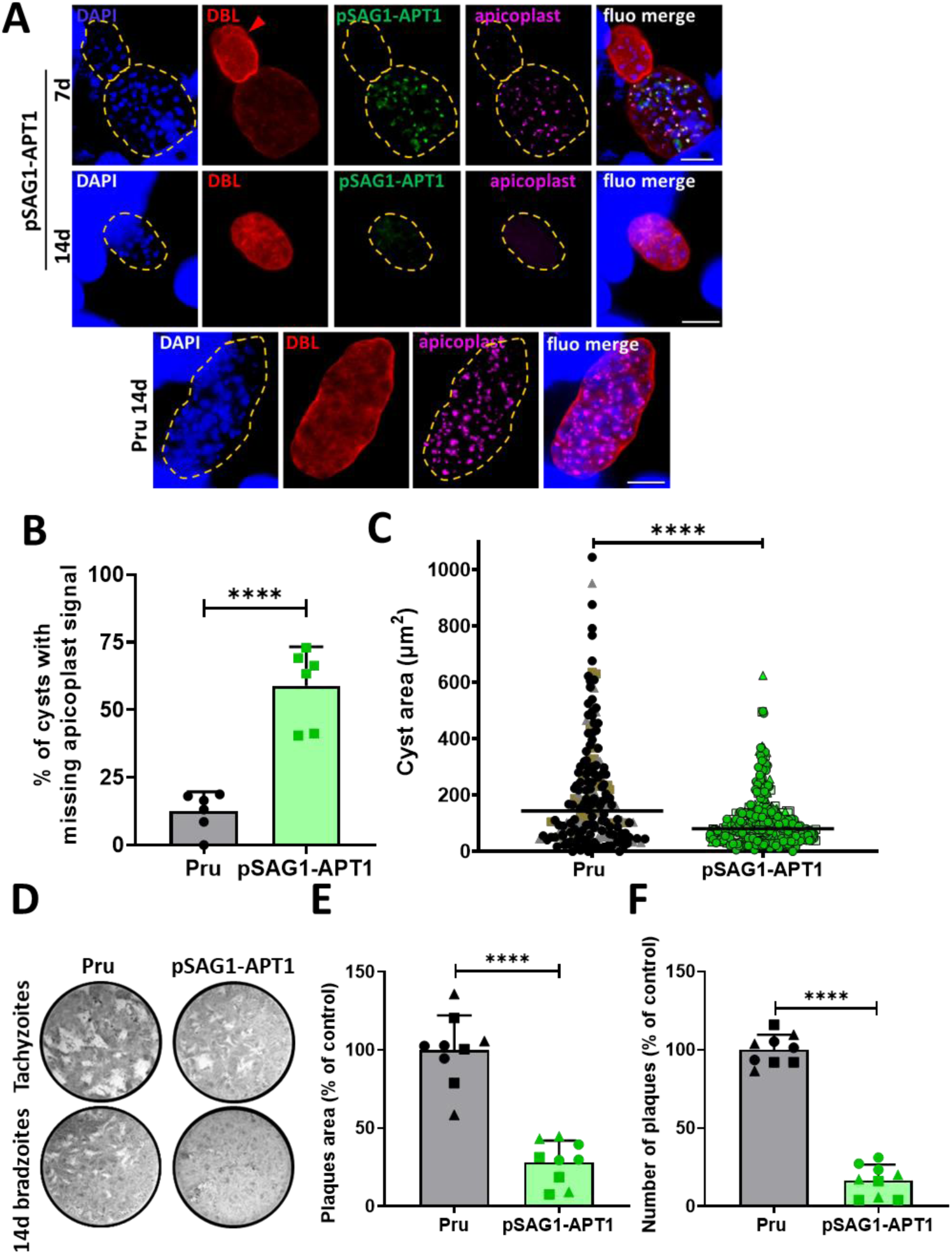
Depletion of APT1 during in vitro differentiation impacts apicoplast maintenance and parasite development and viability. A) Alkaline pH-induced *in vitro* differentiation of pSAG1-APT1 parasites for 7 or 14 days leads to the disappearance of APT1 and of the Cpn60 apicoplast marker (magenta) in mature cysts that are more intensely labelled with DBL (arrowhead). A 14 day-differentiated cyst of the Pru cell line is shown as a control. Cyst wall is outlined with dashed lines. DNA was stained with DAPI. Scale bar=10 µm. B) The integrity of the Cpn60-labelled apicoplast signal was quantified after 14 days of alkaline pH-induced differentiation and showed a significant loss in the pSAG1-APT1 cell line. Data are mean from *n*=6 independent experiments +SD. **** *p* ≤ 0.0001, unpaired Student’s *t*-test. C) Quantification of the area of *in vitro*-differentiated cysts kept for 14 days under alkaline pH stress shows that the pSAG1-APT1 cell line generates smaller cysts. Data are mean from *n*=3 independent experiments. Symbols are matched between identical experimental groups. **** *p* ≤ 0.0001, non-parametric Mann-Whitney test. D) Plaque assays were carried out by infecting HFF monolayers with tachyzoites or bradyzoites obtained after 14 days of alkaline pH stress and FACS-based isolation. GFP-negative bradyzoites of the pSAG1-APT1 cell line generates less and smaller plaques than the Pru control cell lines. Measurements of lysis plaque area E) and plaque number F) highlight a significant defect in the lytic cycle of pSAG1-APT1 bradyzoites. Values are means of nine values (representing *n*=3 independent experiments, containing three technical triplicates) +SD. Symbols are matched between identical experimental groups. Mean value for the Pru cell line was set to 100% as a reference. **** *p* ≤ 0.0001, unpaired Student’s t test.

To assess the impact of APT1 down-regulation on *in vitro* cyst development, we imaged cysts after two weeks of differentiation and then measured cyst area (Fig. 2C). We found the mean area to be significantly lower in the pSAG1-APT1 cell line compared with the parental Pru cell line. This suggests that APT1 depletion upon *in vitro* differentiation has a negative impact on cyst development. We next sought to assess the impact of APT1 depletion on bradyzoite viability using plaque assay as a proxy, as described previously (36). Cysts were generated *in vitro*, bradyzoites were released by pepsin treatment and APT1-negative parasites were isolated with a cell sorter based on the absence of a GFP signal, and seeded on host cell monolayers (Fig. S4A, B). APT1-negative bradyzoites were found to generate many fewer and much smaller plaques than the Pru parental cell line (Fig. 2D, E, F). Of note, by *in vitro* plaque assay tachyzoites of the pSAG1-APT1 cell line did not show any particular fitness defect compared with the Pru parental cell line (Fig. 2D, S5A, B).

In conclusion, our results indicate that specific depletion of APT1 in *in vitro*-generated bradyzoites leads to visible consequences on the homeostasis of the apicoplast and on bradyzoite viability.

### pSAG1-APT1 parasites induce a severe acute phase in mice, yet *ex vivo*-generated bradyzoites are impaired in fitness

While *in vitro*-derived cysts are certainly a useful model, some important differences have been observed with *in vivo*-derived cysts (20), thus we sought to confirm our findings with additional assays in an animal model. To this end we infected CBA mice with the pSAG1-APT1 and Pru control cell lines. In three independent experiments we observed an excessive mortality and a more pronounced body mass loss in the surviving animals for mice infected with pSAG1-APT1 tachyzoites (Fig. 3A, B), hinting that infection with this mutant leads to a severe acute phase. As dendritic cells act early both as immune response mediators and as parasite carriers that facilitate the dissemination of the infection and may play a key role in the acute phase (45), we wanted to evaluate the ability of pSAG1-APT1 tachyzoites to infect these cells *in vitro* using a cytometry based assay (Fig. S5C, D) (46). Yet, our results showed that pSAG1-APT1 tachyzoites do not infect dendritic cells more efficiently than the Pru cell line (Fig. S5E).

**Figure 3.**
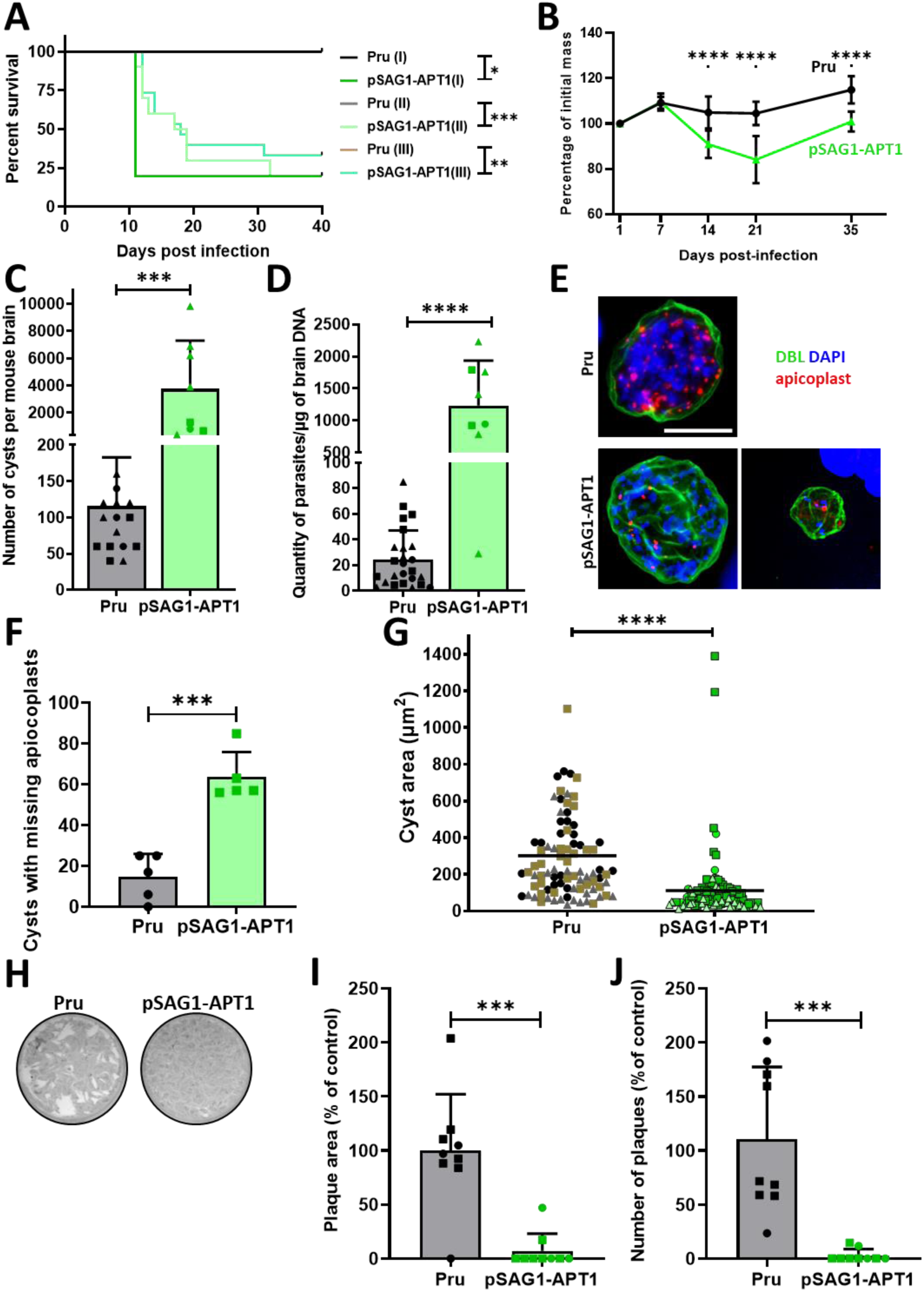
Infection of mice with pSAG1-APT1 parasites leads to a severe acute phase and high cyst burden but generates essentially non-viable bradyzoites. A) Survival curves of CBA mice infected with 100 parasites of the Pru or pSAG1-APT1 cell lines shows that the latter induces an excessive mortality during acute phase. Data from *n*=3 independent experiments are plotted. * *p* ≤ 0.05, ** *p* ≤ 0.01, *** *p* ≤ 0.001, log-rank Mantel-Cox test. B) Monitoring of body mass of mice infected with Pru or pSAG1-APT1 tachyzoites. Infection with pSAG1-APT1 induces a more important mass loss than the control cell line, reflecting a more severe acute phase. Data represent mean from *n*=3 independent experiments +SD. **** *p* ≤ 0.0001, unpaired Student’s t test. C) Quantification of cyst burden in infected mice forty days post-infection. Values are from *n*=3 independent experiments and are expressed as means +SD. Symbols are matched between identical experimental groups. *** *p* ≤ 0.001, unpaired Student’s t test. D) Parasite burden measured by qPCR on genomic DNA from infected mice. Values are from *n*=3 independent experiments and are expressed as means +SD. Symbols are matched between identical experimental groups. **** *p* ≤ 0.0001, unpaired Student’s t test. E) Labelling of mouse brain-derived cysts with anti-Cpn60 shows partial loss of the apicoplast in the pSAG1-APT1 cell line. Scale bar=10 µm. F) Quantification of Cpn60 signal in brain-derived pSAG1-APT1 cysts shows that they display partial or total signal loss 40 days post-infection. Shown are the mean +SD from *n*=5 independent experiments, at least 12 cysts for each condition were assessed. *** *p* ≤ 0.01, unpaired Student’s *t*-test. G) Quantification of the area of brain-derived cysts obtained 40 days post-infection shows that the pSAG1-APT1 cell line generates smaller cysts. Data are mean from *n*=3 independent experiments. Symbols are matched between identical experimental groups. **** *p* ≤ 0.0001, non-parametric Mann-Whitney test. H) A representative plaque assay carried out by infecting HFF monolayers with brain-derived bradyzoites obtained after 40 days of infection shows that the pSAG1-APT1 cell line generates very few plaques. Measurements of lysis plaque area I) and plaque number J) confirms that there is a significant defect in the lytic cycle of brain-derived pSAG1-APT1 bradyzoites. Values are means from nine replicates from *n*=2 independent experiments +SD. Mean value for the Pru cell line was set to 100% as a reference. Symbols are matched between identical experimental groups. *** *p* ≤ 0.001, unpaired Student’s t test.

Tissue cysts were recovered from the brains of infected mice at 40 days post-infection and quantified by microscopic observation after DBL staining and parasite load was also quantified by quantitative PCR (qPCR, Fig. 3C, D). Mice infected with the pSAG1-APT1 cell line had a much higher cyst burden than the Pru parental cell line, which could either be related to the more severe acute phase observed with these parasites, or illustrate more frequent parasite reactivation. Tissue cysts were imaged to detect the presence of an apicoplast, and a majority of pSAG1-APT1 cysts harbored bradyzoites missing apicoplast-resident proteins (Fig. 3E, F, Fig. S2D, E). Using spectral flow cytometry to reduce the autofluorescence background we initially observed in *in vitro*-differentiated bradyzoites (Fig. S4A, B), we could see that very few APT1-GFP-positive bradyzoites remain after *in vitro* or *in vivo* differentiation, demonstrating an efficient down-regulation of APT1 (Fig. S4C, D, E). We also found that the size of pSAG1-APT1 cysts was smaller than the size of cysts from the Pru cell line (Fig. 3G). Moreover, to assess the viability of these bradyzoites, we performed plaque assays with parasites recovered from these *ex vivo* cysts and observed much less and much smaller plaques with pSAG1-APT1 parasites than with the Pru parental cell line (Fig. 3H, I, J). Finally, parasites recovered from *in vivo* cysts were left for 3 hours to invade HFFs and checked by IFA for the expression of APT1-GFP or the Cpn60 apicoplast marker (Fig. SA, B, C) and while very few were still expressing the APT1 signal (in line with the spectral cytometry quantifications), all were found to retain the Cpn60 signal. Importantly, this suggests that while almost all bradyzoites lost the expression of APT1, only the few not displaying a severe perturbation of the apicoplast were viable enough to invade, highlighting the importance of the organelle for bradyzoite fitness.

In conclusion, while in an *in vivo* setting, pSAG1-APT1 tachyzoites induce a more severe acute phase and lead to a higher parasitemia in the brain, the phenotype of brain-derived pSAG1-APT1 parasites regarding cyst size, loss of the apicoplast or overall viability was very similar to the results we obtained with *in vitro*-generated parasites. Our data thus indicate that the specific loss of APT1 function in *in vitro*- or *in vivo*-derived bradyzoites affects apicoplast homeostasis, bradyzoite development and viability. This suggests that the apicoplast plays an important role for the fitness of bradyzoites.

### Generating a structural mutant of the apicoplast through a dominant-negative approach

Although the exact nature of the metabolic requirement of bradyzoites is unknown, APT1 likely has an important role in the metabolic contribution of the apicoplast for the development of these parasite stages. Yet, as the inhibition of metabolic versus housekeeping functions of the organelle can lead to different phenotypic outcomes (47), we sought to generate a second apicoplast mutant that would be affected in its ability to divide rather than to perform a particular metabolic function. To this end we targeted DrpA, a dynamin-related protein that was previously shown to be essential for apicoplast fission and inheritance in dividing tachyzoites (Fig. 4A) (48). A single mutation in the sequence of the DrpA protein (changing a lysine to an alanine in the GTP-binding site) has been shown to disrupt dynamin function in a dominant-negative fashion in *T. gondii* tachyzoites (48).

**Figure 4.**
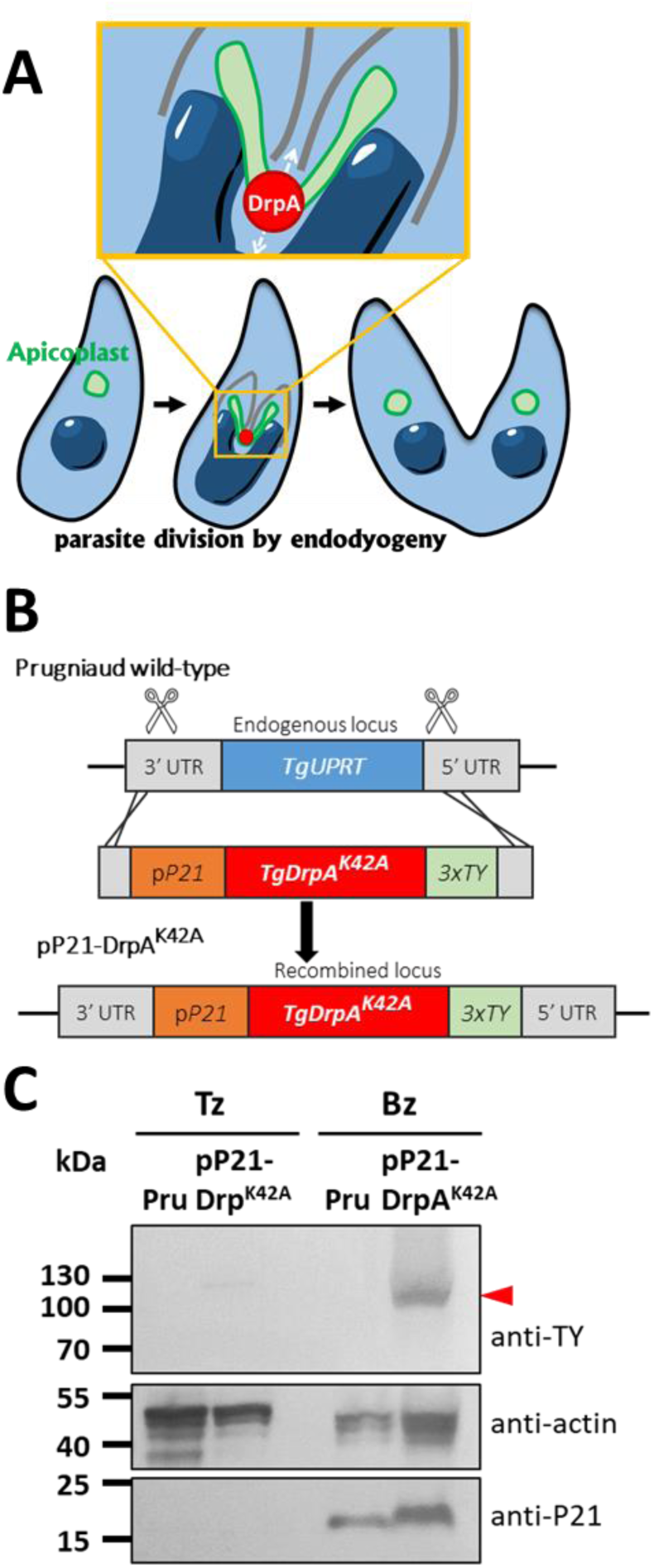
Generating a parasite line expressing a bradyzoite-specific dominant-negative DrpA mutant. A) Schematic representation of DrpA function in apicoplast fission and inheritance during parasite division. B) Scheme describing the strategy to edit the *UPRT* locus by homologous recombination, in order to introduce a mutated version of the *DrpA* gene and express the DrpA^K42A^ protein under the dependence of a bradyzoite-specific *P21* promoter. C) Immunoblot analysis of protein extracts from tachyzoites (Tz) and 6 days *in vitro*-differentiated (apicidin treatment) bradyzoites (Bz) with an anti-TY antibody to detect the expression of the DrpA^K42A^ protein (arrowhead) under the dependence of the *P21* promoter. Actin was used as a loading control and P21 as a differentiation control.

We thus designed a strategy to specifically express the mutant version of DrpA (DrpA^K42A^) in bradyzoites, by using CRISPR/Cas9 to insert the sequence coding for DrpA^K42A^ at the *uracil phosphoribosyl transferase* (*UPRT*) locus (Fig. 4B), which is dispensable for bradyzoites in normal culture conditions and for cystogenesis *in vivo* (49), but allows selection of transgenic parasites with fluorodeoxyribose (50). To drive the expression of the dominant-negative version of DrpA selectively in bradyzoites, we chose the bradyzoite-specific promoter of P21, a late marker of bradyzoite differentiation (51) that is suitable for this type of strategy (52). Our construct also allowed the addition of a TY epitope tag (53) for subsequent immunodetection of the mutated DrpA version. Transgenic parasites, named pP21-DrpA^K42A^, were verified by PCR (Fig. S7) and used for *in vitro* bradyzoite conversion. The UPRT is involved in pyrimidine salvage, which can be compensated by *de novo* synthesis in the parasite (54), but this synthesis relies on CO_2_, which has to be omitted from the classical alkaline pH stress differentiation protocol. We thus used treatment with the histone deacetylase inhibitor apicidin (55, 56) as an alternative for initiating *in vitro* bradyzoite conversion of the pP21-DrpA^K42A^ cell line. Immunoblot analysis showed that the TY-tagged DrpA^K42A^ protein is specifically expressed in *in vitro*-differentiated bradyzoites (Fig. 4C). This was confirmed by immunofluorescence microscopy (Fig. 5A), showing that the DrpA^K42A^ protein was essentially absent from tachyzoites, while it was expressed in differentiated parasites. There, it localized to the cytoplasm and to discrete sub-compartments of the apicoplast as expected (Fig. 5A) (48).

**Figure 5.**
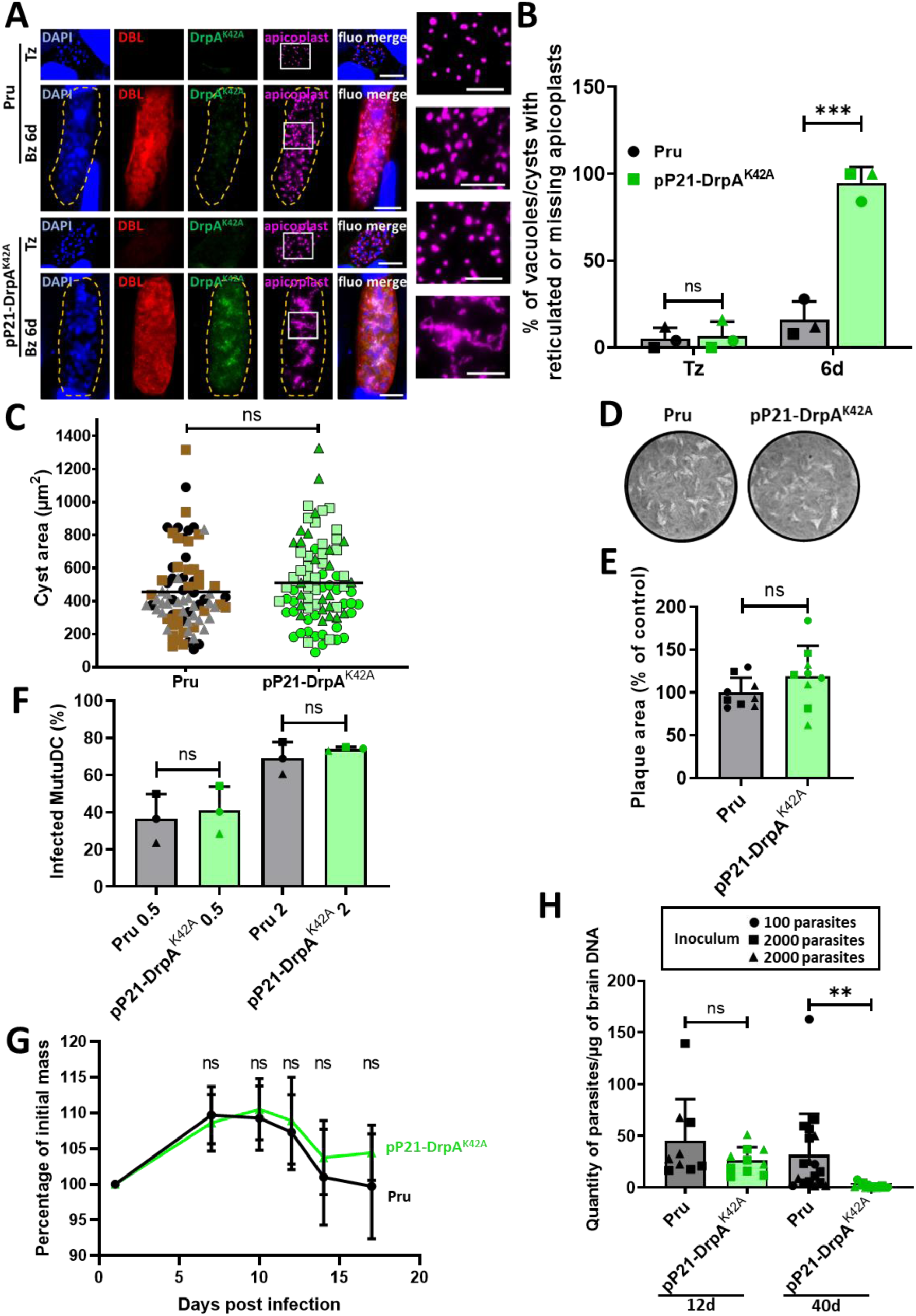
DrpA^K42A^-expressing bradyzoites are impacted for apicoplast division and fail to establish chronic phase in mice. A) Upon apicidin-induced *in vitro* differentiation for 6 days, the DrpA^K42A^ protein is expressed and bradyzoites show extensive reticulation of their apicoplast (Cpn60-labelled, magenta, magnified in insets) in DBL-labelled cysts. The Pru parental cell line and tachyzoites of the two cell lines are shown as a control. Cyst wall is outlined with dashed lines. DNA was stained with DAPI. Scale bar=10 µm (5 µm for insets). B) The proportion of cysts with abnormal apicoplast signal (reticulated or lost) after 6 days of apicidin treatment was quantified. Values are means from *n*=3 independent experiments +SD. Symbols are matched between identical experimental groups. *** *p* ≤ 0.001, ‘ns’ not significant, unpaired Student’s t test. C) Quantification of the size of *in vitro*-differentiated cysts kept for 6 days in the presence of apicidin shows no size difference between cysts of the pP21-DrpA^K42A^ cell line and the Pru control. Data are mean from *n*=3 independent experiments. Symbols are matched between identical experimental groups. ‘ns’ not significant, non-parametric Mann-Whitney test. D) A representative plaque assay carried out by infecting HFF monolayers with tachyzoites shows that the pP21-DrpA^K42A^ parasites are not affected in their lytic cycle *in vitro*. E) Measurement of lysis plaque areas confirms that there is no major defect in the lytic cycle of pP21-DrpA^K42A^ tachyzoites. Values are means from *n*=3 independent experiments (each one with three technical replicates) +SD. Mean value for the Pru cell line was set to 100% as a reference. Symbols are matched between identical experimental groups. **** *p* ≤ 0.0001, unpaired Student’s t test. F) *In vitro* dendritic cell infection assay was performed and analyzed by flow cytometry and shows no impairment of the invasive capacity of pP21-DrpA^K42A^ tachyzoites. Data are mean from *n*=3 independent experiments; ‘ns’ not significant, unpaired Student’s t test. G) Monitoring of body mass of mice infected with Pru or of pP21-DrpA^K42A^ tachyzoites shows that the acute phase occurs normally with of pP21-DrpA^K42A^ parasites. Data represent mean from 20 individual mice from *n*=2 independent experiments +SD. ‘ns’ not significant, unpaired Student’s t test. H) Parasite burden at 12 days or 40 days post-infection measured by qPCR on genomic DNA from infected mice. Values are from *n*=2 (12 days timepoint) or *n*=3 (40 days timepoint) independent experiments, performed with an inoculum of 100 parasites or 2000 parasites, and are expressed as means +SD. ** *p* ≤ 0.01, ‘ns’ not significant, unpaired Student’s t test.

### Dominant-negative pP21-DrpA^K42A^ parasites have their apicoplast specifically affected at the bradyzoite stage but show no fitness defect as tachyzoites

Upon differentiation using apicidin, detailed analysis of DrpA^K42A^-expressing bradyzoites revealed that they display a reticulated apicoplast labelling or absence of the organelle (Fig. 5A, B, S8), which is similar to what was previously described upon DrpA^K42A^ expression in tachyzoites (48). Apicidin triggered stage conversion very efficiently, but long-term incubation may have unspecific effects, so we also used alternative methods of differentiation to drive DrpA^K42A^ expression and confirmed apicoplast segregation and reticulation problems within cysts upon short term alkaline stress (as mentioned above long term treatment is detrimental in absence of the *UPRT* locus) and heat stress-induced differentiation (57) (Fig. S8A). Strikingly, when using an anti-P21 antibody on differentiating pP21-DrpA^K42A^ parasites, we could see that cysts/vacuoles in which the morphological impact on the apicoplast was the most visible were those with a higher expression of P21, confirming that the dominant-negative effect is efficiently regulated by the bradyzoite-specific promoter (Fig. S8B). However, when cyst size was measured after apicidin-induced *in vitro* differentiation, in contrast to the results obtained with pSAG1-APT1 parasites, no significant difference was found in comparison with the control (Fig. 5C). This highlights potential differences on the timing of phenotype initiation or on the consequences at the cellular level when affecting the metabolism versus the ultrastructure of the apicoplast. This also called for long term assessment of viability the pP21-DrpA^K42A^ parasites in the mouse model. Prior to implementing these *in vivo* experiments, we assessed the *in vitro* fitness of the tachyzoite stage of the pP21-DrpA^K42A^ cell line and found no particular problem in their ability to form plaques on fibroblast monolayers (Fig. 5D, E). Like described earlier we also checked by flow cytometry analysis that the pP21-DrpA^K42A^ tachyzoites were capable of infecting dendritic cells (46).Our results showed that pP21-DrpA^K42A^ tachyzoites infect dendritic cells as efficiently as the Pru cell line (Fig. 5F).

pP21-DrpA^K42A^ tachyzoites, when injected into mice, induced mass loss in the animals in the timeframe corresponding to the acute phase of infection (i.e. up to 17 days, Fig. 5G), hinting that this part of the infection process was occurring normally. However, in contrast to the pSAG1-APT1 cell line, infection with pP21-DrpA^K42A^ tachyzoites did not induce any specific mortality in mice during the acute phase (Fig. S9A). Parasite quantification by qPCR showed the presence of parasites in the brain as early as 12 days post-infection, confirming that pP21-DrpA^K42A^ mutant parasites are able to disseminate and invade the brain; yet, very little pP21-DrpA^K42A^ parasites were detected 40 days post-infection (Fig. 5H). Importantly, there was no difference in the outcome at 40 days when we increased the inoculum from 100 to 2000 tachyzoites (Fig. 5H). In fact, 40 days post-infection, brain samples from mice infected with the pP21-DrpA^K42A^ cell line contained either very few or no detectable cysts, as quantified by qPCR or estimated by microscopic counts (Fig. 5H, Fig. S9B). The fitness or invasive capacity of pP21-DrpA^K42A^ tachyzoites is not apparently affected *in vitro* (Fig. 5D, E, F) and our *in vivo* data shows that they likely generate a normal acute phase and are able to reach the brain (Fig. 5G, H), however long-term persistence is affected as illustrated by the almost complete absence of detectable parasites 40 days post infection (Fig. 5H, Fig. S9B). This suggests that long term perturbation of apicoplast division impacts bradyzoite viability and persistence *in vivo*.

## Discussion

The long-term persistence of encysted *T. gondii* parasites in a large proportion of the human population, particularly in ocular tissue and in the central nervous system, has potentially important consequences on the development of a future pathology. As neither the immune system nor current drug-based approaches seem able to eliminate this developmental stage (31, 58), it is essential to look for novel therapeutic approaches. Because of its evolutionary history and its metabolic importance, the apicoplast has long been considered a promising drug target in the tachyzoite stage, in which both pharmacological and genetic studies have shown that loss of the organelle, loss of its genome, or loss of its metabolic function result in the death of the parasites. Here, for the first time, we used two independent and complementary genetic approaches to investigate the importance of the apicoplast in the encysted bradyzoite stage.

While there has been recent progress in the *in vitro* methods used to initiate stage conversion into bradyzoites (18–20), the complexity and diversity of bradyzoites may not always be fully recapitulated *in vitro*, so the most canonical way to experimentally generate authentic mature *T. gondii* tissue cysts is currently through murine infections. Regulating gene expression in a cyst-enclosed stage that is mostly confined to neuronal tissues (whose access is restricted by the blood-brain barrier), is a technical challenge as it may not be reached efficiently by compounds commonly used for driving conditional silencing. For this reason, we chose a stage-specific promoter-driven strategy to conditionally control gene expression. Although this does not allow complete control of the timing of protein expression, we and others have successfully used the *SAG1* promoter swap strategy to achieve selective loss of expression of a gene of interest in bradyzoites in the past (36, 59, 60). We also took advantage of stage-specific expression in a complementary approach, this time with the promoter of the late bradyzoite marker P21 (52), to specifically switch on the expression of the dominant-negative mutant version of the DrpA protein. Our data show that the regulation of expression was efficient, both for *in vitro* and *in vivo* bradyzoite conversion models. However, quantifications of the effect on the apicoplast morphology or the loss of the organelle, while highlighting a strong impact on the organelle, also showed occasionally some phenotypic differences. For instance, apicoplasts were still present in a few cyst-enclosed pSAG1-APT1 bradyzoites recovered 40 days post infection (Fig. 3E, Fig. S2D, Fig. S6), in spite of a very efficient down-regulation of APT1 (Fig. S4C, D, E, Fig. S6). Although these APT1-deficient organelles may already not be fully functional, it seems that a stronger effect on the apicoplast occurs later and with some degree of heterogeneity. This is not surprising because stage conversion is a continuum that may happen asynchronously. As a result, there can be some heterogeneity in gene expression, as single-cell RNAseq analyses of *in vitro*-differentiated parasites suggest (61). Analyses on brain-derived cysts also highlighted a fairly large diversity between cysts and among bradyzoites within the same cyst (5, 62). Our work nevertheless confirms that stage-specific promoters can be used efficiently to investigate gene function in bradyzoites through a simple promoter replacement.

We purposely used different approaches and different types of mutants affected in apicoplast homeostasis, with one primarily impacted for metabolism and the other one in organelle fission and inheritance: we wanted to see if there was converging evidence on the essentiality of the organelle in bradyzoites. Interestingly, the two cell lines we generated through these different approaches were found to display phenotypes that are not completely overlapping (Fig. S10). One particularly striking difference is the strong acute phase induced by the pSAG1-APT1 cell line (Fig. 3A, B, C and D). One possible explanation is that promoter replacement likely led to an overexpression of APT1 (Fig. S1C and D), and this might in turn have conferred a metabolic and thus a fitness advantage to tachyzoites in the host. On the other hand, our plaque assays show that there does not seem to be an increased fitness for tachyzoites *in vitro* (Fig. 2D, S3). Discrepancies between *in vitro* and *in vivo* settings have been highlighted in several studies describing the metabolic flexibility of *T. gondii* tachyzoites, which are able to scavenge a number of nutrients in rich culture medium but may encounter more restrictive conditions in the animal host (63). A number of host and parasite factors account for the heterogeneity in the ranges of mice brain cyst burden described in the literature, and while it is difficult to establish an exact correlation between the cyst burden and the initial inoculum, it is quite likely that a more successful acute phase can lead to increased parasite load in the brain (64, 65). It is thus possible that a fitness advantage during the acute phase would be the reason of the high cyst numbers generated by the pSAG1-APT1 cell line. These large cyst numbers may also potentially reflect multiple reactivation and reinvasion events, which might also explain the fact that these cysts were generally found to be smaller than those of the parental cell line. However, the majority of pSAG1-APT1 cysts that were recovered showed morphological alterations, especially regarding the apicoplast (Fig. 3E and D, Fig. S2). Moreover, bradyzoites that were released from these cysts were found to be largely non-viable (Fig. 3H, I and J). So, altogether our data rather suggest that the small cysts reflect a problem in parasite development instead of recent parasite establishment. Importantly, no *in vivo*-generated bradyzoite with a strong apicoplast defect (i.e. lacking apicoplast marker labelling) was fit enough to invade host cells upon release from cysts (Fig. S6), further supporting the importance of the organelle for bradyzoite fitness.

In contrast, the pP21-DrpA^K42A^ cell line largely failed to generate cysts (Fig. 5H). We could establish that it was not due to an alteration in the acute phase, as the parasites were found to reach the brain at the end of this phase of infection (Fig. 5G and H). Instead, this probably rather highlights an increased elimination by the immune system, or/and a strong defect in cystogenesis. While both our strategies were aimed at interfering with the function of the apicoplast, the different outcomes in the animal model concerning cyst generation may be due to the different promoters that were used, which respectively turn genes on or off at different times during the stage conversion process, thus affecting the timeframe of the manifestation of the detrimental phenotypes.

Another main difference in the mutants we generated is that while for both there was a visible morphological impact *in vitro*, the pSAG1-APT1 mutant cell line was already strongly affected in cyst growth during *in vitro* differentiation (Fig. 2C), while for pP21-DrpA^K42A^ bradyzoites the detrimental effect on cystogenesis only manifested during long term infection in the animal (Fig. 5C and H). This could be related to the fact that the former targets metabolic pathways hosted by the organelle, while the latter targets its division and inheritance in parasite progeny. It has been known for some time that there are differences in the kinetics of parasite demise whether a metabolic pathway of the apicoplast or a housekeeping function of the organelle are targeted (47). Targeting apicoplast-hosted metabolism through genetic or pharmaceutical-based approaches usually lead to a rapid death of the parasites, while affecting apicoplast division usually leads to a so-called delayed death, where parasites are only affected after another round of reinvasion. While *in vitro*-differentiated pP21-DrpA^K42A^ bradyzoites showed marked signs of morphological alteration of their apicoplasts (Fig. 5A, Fig. S8), the remaining organelles potentially retained some metabolic capacity. A strong impact on the apicoplast, which is linked to organelle inheritance defects in this mutant, likely depend on the parasites undergoing several successive rounds of division that would only happen over a longer period of time. It was shown that *T. gondii* tachyzoites can potentially share resources (metabolites and proteins) through a connection at their basal pole, ensuring proper cellular division in spite of an absence of apicoplast of several parasites within the PV, and thus explaining the delayed death effect (66). However, bradyzoite division is largely asynchronous and there is no connection between the parasites in mature cysts (9, 66, 67), which suggests that organelle loss may be more detrimental for this developmental stage. Moreover, this might be particularly important in an *in vivo* setting, where nutrient sources are potentially scarce, which could explain why cystogenesis of pP21-DrpA^K42A^ parasites ended up being strongly affected in these conditions.

Importantly, in the mouse model, both pSAG1-APT1 parasites (that generate numerous but small cysts, containing less fit bradyzoites) and pP21-DrpA^K42A^ parasites (that display a strong defect in cystogenesis) showed converging evidence that the apicoplast is important for *T. gondii* survival and persistence as bradyzoites. This now raises the question of which essential function(s) the organelle is performing for this developmental stage. An earlier *in vivo* morphological study showed that duplication or elongation of the apicoplast could still be observed in bradyzoites many weeks or even months post-infection, suggesting an active role for the organelle (68). Due to the lack of functional studies, the metabolic activities and requirements of bradyzoites are, however, still largely uncharacterized. RNA expression data from *in vivo*-generated cysts suggest that the four main metabolic pathways hosted by the apicoplast could be active in bradyzoites, with enzymes of the iron-sulfur cluster and isoprenoid synthesis pathways even potentially upregulated (16). It would now be particularly interesting to apply our conditional strategies to specifically interfere with individual enzymes of these pathways in bradyzoites.

While finely-tuned control of protein expression is technically difficult to achieve directly in brain-encysted mature bradyzoites, through this work we have provided corroborating lines of evidence showing that the organelle is important for bradyzoite fitness and persistence. Importantly, this validates the organelle as a potential drug target in the context of the establishment of chronic toxoplasmosis. Of course, it should be kept in mind that designing drugs able to interfere with the apicoplast in bradyzoites will constitute a major challenge. For instance, long-term treatment in mice suggests that apicoplast-targeting drugs like clindamycin and spiramycin can both reduce cyst burden, yet they may not completely eradicate the parasites (69, 70), and novel compounds targeting a specific pathway hosted by the organelle would have to be able to go through a series of physical hurdles that include the blood-brain barrier, the host cell plasma membrane, the cyst wall, the parasite plasma membrane and, finally, the four membranes of the organelle. Also, as bradyzoites are slow growing, they are likely to be less sensitive to metabolic targeting, by analogy with bacterial or tumoral ‘persister’ cells that are notoriously prone to resist to drugs (71). However, although cyst-enclosed bradyzoites display some heterogeneity in their cellular and metabolic activities, they remain dynamic growing entities (62), and our *in vivo* results suggest that they are likely to be vulnerable to long-term interference with apicoplast-related functions. So, the possibility to exploit this as a therapeutic strategy should not be overlooked.

## Materials and Methods

### Animal care and ethics statement

Animal care and use protocols were carried out under the control of the National Veterinary Services and in accordance with the European Union guidelines for the handling of laboratory animals EEC Council Directive, 2010/63/EU, September 2010). The protocol inducing pain (APAFIS#25130-2020040721346790 v3) was approved by the local Ethical Committee for Animal Experimentation registered by the ‘Comité National de Réflexion Ethique sur l’Expérimentation Animale’ under no. CEEA122. CBA/J mice mice were purchased from Janvier Labs (France).

### Parasites and cells culture

Tachyzoites of the Prugniaud *T. gondii* strain (72), as well as derived transgenic parasites, were maintained by serial passage in monolayers of human foreskin fibroblast (HFF, American Type Culture Collection, CRL 1634) grown in Dulbecco’s modified Eagle medium (DMEM, Gibco), supplemented with 5% of decomplemented fetal bovine serum, 2 mM glutamine, and a cocktail of penicillin-streptomycin at 100 μg/ml.

### *In vitro* differentiation

*In vitro* conversion into bradyzoites was achieved either through alkaline pH stress (57), or by treating with the histone deacetylase inhibitor apicidin (55), or by heat-stress (57).

For alkaline pH-induced differentiation (57) mostly used for conversion of the pSAG1-APT1 cell line, HFF monolayers of cells grown were infected with freshly egressed tachyzoites for 24h in DMEM culture medium, either on T25 flasks (with 500,000 parasites) or on coverslips in 24-well plates (with 50,000 parasites). The medium was then replaced by Minimum Essential Medium (MEM; 10X) without NaHCO3 (Gibco), supplemented with 50 mM HEPES, 1% penicillin-streptomycin, 1% Glutamine and 3% FBS and adjusted to pH 8.2. Parasites were then cultured at 37°C without CO_2_ the medium was changed every two days during the whole duration of the differentiation experiment, which lasted up to two weeks.

For the pP21-DrpA^K42A^ cell line that cannot sustain long-term CO_2_ deprivation, differentiation was induced by alkaline pH for only 3 days, or alternatively by treatment with the histone deacetylase inhibitor apicidin (55) or by heat-stress (57). Briefly, for apicidin-induced differentiation tachyzoites were used to infect host HFFs and after 24h they were treated with 40 nM apicidin (Santa Cruz) for up to 6 days. It should be noted that longer incubations with apicidin led to some parasite mortality, even in the control cell line. Alternatively, stage conversion in the pP21-DrpA^K42A^ cell line was initiated by a temperature shift. Briefly, host cells were put at 43°C for two hours prior to infection to acquire thermotolerance and returned at 37°C for an hour. Then tachyzoites were allowed to invade for 2h before placing the culture at 43°C for two days and then returned at 37°C for up to six days.

### Generation of the TgAPT1 conditional knockdown pSAG1-APT1 cell line

To generate a conditional knock-down TgAPT1 line specifically depleted at the bradyzoite stage, the endogenous promoter of the *TgAPT1* gene was replaced by a *SAG1* promoter (a tachyzoite-specific promoter) using a CRISPR/Cas9-based strategy. The whole genomic sequence of TgAPT1 (www.toxodb.org accession number TGME49_261070) was amplified by PCR with the Q5 High Fidelity polymerase (New England Biolabs) from Pru genomic DNA, with primers ML3012/ML3013 (all the primers used in this study are listed in Table S1). The resulting 1059 bp fragment was then inserted into the pLIC-CAT-GFP plasmid (74), in frame with the sequence coding for the GFP, resulting in the pLIC-TgAPT1-GFP plasmid. The DHFR-SAG1 promoter cassette was amplified from the DHFR-pSAG1-GFP-TgATG8 plasmid (36) using primers ML3015/ML3016, both containing PacI restriction site. The DHFR-SAG1 promoter cassette was then digested and inserted into PacI-digested pLIC-TgAPT1-GFP plasmid to create DHFR-pSAG1-TgAPT1-GFP plasmid.

Specific single-guide RNAs (sgRNAs) were generated to introduce a double-stranded break at the 5’ and 3′ boundaries of the *TgAPT1* locus. Primers used to generate the guides were ML3026/ML3027 and ML3030/ML3031, and the protospacer sequences were introduced in the Cas9-expressing pU6-Universal plasmid (Addgene, ref #52694) (75). The cassette corresponding the DHFR-pSAG1-TgAPT1-GFP cassette was amplified by PCR from the plasmid of the same name using primers ML3088/ML3089 with the KOD DNA polymerase (Sigma-Aldrich). This donor sequence was then used to transfected to Pru background, together with the two guide-expressing plasmids. The Electro Cell Manipulator 630 (BTX) was used with the following settings: 2.02 kV, 50 Ω, 25µF. Transfected parasites were selected with 1µM pyrimethamine and cloned by limit dilution. Positive parasites were first screened for GFP-positive apicoplasts by microscopy, then positive clones were verified by PCR using primers ML3023/ML3116 (5’ end of the wild-type locus), ML3115/ML4541 (3’ end of the wild-type locus), ML2873/ML3116 (5’ end after integration) and ML4525/ML4541 (3’ end after integration integration). The resulting parasite line was designated pSAG1-APT1. Several clones were isolated and tested for regulation of protein expression and impact on the apicoplast during *in vitro* differentiation and showed no particular difference, so one was subsequently chosen for complete phenotypic analysis.

### Generation of the conditional dominant-negative pP21-DrpA^K42A^ cell line

The pP21-DrpA^K42A^ mutant cell line was generated based on the *pUPRT-*pTUB*-G13-TY* plasmid (76) using a CRISPR-based strategy. We performed PCR with the Q5 High Fidelity polymerase (New England Biolabs) to amplify a 1,500 bp region upstream of the start codon of the *TgP21* gene (www.toxodb.org accession number TGME49_238440 - Me49 being the reference for cystogenic *T. gondii* strains) containing a promoter sequence, using primers ML3655/ML3656 and cloned using NotI/BclI sites in the *pUPRT-*pTUB*-G13-TY* plasmid to create the *pUPRT-*pP21*-G13-TY plasmid. To amplify the* DrpA *coding sequence (*www.toxodb.org accession number #TGME49_267800), total cDNA of the Pru strain was generated following purification of parasite total RNA using the NucleoSpin RNA kit (Macherey Nagel), followed by a reverse transcription using the Super Script III first strand synthesis supermix kit (Invitrogen). This was then used to amplify the *DrpA* cDNA by PCR with the Q5 polymerase (New England BioLabs) and primers ML 4417/4418. *DrpA* cDNA was then subcloned in the PCR-Blunt II-TOPO vector (Invitrogen). Site-directed mutagenesis was performed with the QuikChange II Site-directed Mutagenesis kit (Agilent) and primers ML 4421/4422 to generate the TOPO-DrpA^K42A^ vector. The fragment was then cloned into the BglII/AvrII digested *pUPRT-*pP21*-G13-TY* plasmid in order to create the pUPRT-pP21-DrpA^K42A^-Ty plasmid. This plasmid was then linearized with the AvrII enzyme and dephosphorylated with Antartic Phosphatase (New England Biolabs) in order to insert an extra double TY tag. The sequence coding for the tag was created by annealing primers ML4419/4420 and cloned into the linearized and dephosphorylated pUPRT-pP21-DrpA^K42A^-TY vector, to yield the pUPRT-pP21-DrpA^K42A^-3xTY plasmid.

This plasmid was then linearized by KpnI/NsiI prior to transfection. The recombination efficiency was increased by co-transfecting with Cas9-expressing pU6-UPRT plasmids, generated by integrating *UPRT*-specific protospacer sequences (with primers ML3445/ML3446 for the 5’ and ML2087/ML2088 for the 3’) which were designed to allow a double-strand break at the *UPRT* locus. Transgenic parasites were selected with 10 µM 5-fluorodeoxyuridine (Sigma-Aldrich) and then cloned by serial limiting dilution in a p96 wells plate. The resulting cell line was designated pP21-DrpA^K42A^. Several clones were isolated and tested for regulation of protein expression and impact on the apicoplast during *in vitro* differentiation and showed no particular difference, so one was subsequently chosen for complete phenotypic analysis.

### Mouse infection

Seven-week-old male CBA/J mice (Janvier Labs) were infected intraperitoneally with 100 or 2000 tachyzoites of the appropriate strains. At different timepoints post-infection, the brains were collected and cysts were isolated by isopycnic centrifugation (73), bradyzoite were recovered after pepsin treatment, and parasite quantification was performed as described previously (60).

### Cyst purification and bradyzoite isolation

Cysts were isolated by isopycnic centrifugation (73). Briefly, mouse brains were collected 40 days post infection and homogenized in a Dounce tissue grinder in 3 ml of PBS. To separate parasite cysts from brain cell debris, the homogenate was centrifuged and the pellet was resuspended in 2.5 ml of isotonic Percoll (90% Percoll, 10% NaCl 1.5 M), and PBS was added up to 7.5 ml. After centrifugation at 500 g for 15 minutes at 4°C, cysts were recovered in the pellet for further analysis.

Bradyzoites were recovered from tissue cysts by pepsin treatment. Briefly, tissue cysts were washed in Hanks’ Balanced Salt Solution (HBSS) and incubated with a freshly prepared 2X pepsin solution (340 mM NaCl,120 mM HCl, 0.52 mg/ml pepsin from porcine stomach mucosa – Sigma-Aldrich), and then incubated at 37°C form 30 minutes before stopping the reaction by adding complete DMEM.

*In vitro*-derived bradyzoites of the pSAG1-APT1 or Pru cell lines were removed with a cell scrapper and subjected to a similar treatment, then passed through a 25G needle and a 40 µm filter prior to cell sorting with a FACSCanto flow cytometer (Becton Dickinson). Fluorescence of *in vitro*- or *ex vivo*-derived bradyzoites was also measured with an Aurora cytometer (Cytek), as this spectral system allows autofluorescence extraction for a greater population resolution in the samples (77).

### Mouse infection, quantification of parasite burden by cyst enumeration and qPCR

Seven-week-old male CBA/J mice (Janvier Labs) were housed under specific pathogen-free conditions and infected with 100 or 2000 tachyzoites of the appropriate strains intraperitoneally. Infection was confirmed by assessing seropositivity of 21 days post infection using dot blot analysis with parasite extracts. Mouse health status and body mass were monitored daily throughout acute stage, and for up to 40 days.

At different timepoints post-infection, the brains were collected and homogenized with a glass Potter in PBS. For parasite genomic DNA assessment, gDNA was extracted using the DNEasy Blood & Tissue Kit (Qiagen ref 69506). As described earlier (78), a 529-bp repeat element in the *T. gondii* genome was amplified using the TOX9 and TOX11 primers. The number of parasite equivalent per μg of brain DNA was estimated by comparison with a standard curve, established with a known number of Pru tachyzoites. For cyst enumeration, 5% of the homogenate was stained with fluorescein-conjugated *Dolichos biflorus* agglutinin (Vector Laboratories). Cysts were counted using an inverted fluorescence microscope.

Cyst area from both *in vivo* and *in vitro*-derived cysts was measured based on this cyst wall staining using the ZEN software v2.5 (Zeiss).

### Dendritic cell infection assay

MutuDC, a C57BL/6-derived dendritic cell line obtained from H. Acha-Orbea (46), were plated in a 6-well plate at 5.10^5^ cells per well. Three hours later, MutuDC were infected with either parental Pru, the pSAG1-APT1 line, or the pP21-DrpA^K42A^ line. After 24h, cells were incubated with FcR block and eFluor 450 Fixable Viability Dye (1/1000, eBioscience) in PBS, fixed with 4% PFA, permeabilized in PBS BSA 0,2% Saponine 0,05%, and labeled in that same buffer with AF647-coupled rabbit antibodies that recognize the GRA6 C-terminal decapeptide HPGSVNEFDF (custom-made, Biotem, Grenoble) for 1h at 4°C, allowing to detect vacuoles in which GRA6 was actively secreted by live parasites. Samples were post-fixed using 1% PFA before acquisition on a BD Fortessa and analyzed using FlowJo software.

### Plaque assays

For plaque assays, confluent monolayers of HFFs seeded in 24 well plates were infected with freshly egresses tachyzoites or freshly isolated bradyzoites. Parasites were seeded in the first lane of the plate and diluted by 1/3 in each subsequent row. The number of parasites used in the first wells was as follows: 7000 (tachyzoites, manually counted), 3000 (FACS-sorted bradyzoites), or up to 4000 (qPCR-estimated tissue cyst-derived bradyzoites). Parasites which were left to grow for 10 days for tachyzoites and up to 20 days for bradyzoites. They were then fixed with 4% v/v PFA and plaques were revealed by staining with a 0.1% crystal violet solution (V5265, Sigma-Aldrich). Pictures of the plaques were acquired with an Olympus MVX10 microscope and plaques area were measured using the ZEN software v2.5 (Zeiss). Plaque area and plaque number of the mutant cell lines were expressed relatively to the mean value of those generated by the Pru control strain that was set to 100%

### Immunofluorescence microscopy

Immunofluorescence assays (IFAs) of intracellular parasites were performed on coverslips containing infected HFF monolayers, fixed with 4% paraformaldehyde in PBS for 20 minutes at room temperature. Cells were washed three times with PBS, permeabilized with 0.3% Triton X-100/PBS for 15 minutes and then saturated with a 1% w/v bovine serum albumin (BSA)/PBS blocking solution for 30 minutes. Proteins were stained with primary antibodies for 1h, followed by three washes with PBS before incubation with secondary antibodies in a 1% BSA/PBS solution for 1h. Primary antibodies used in this study and their respective dilutions were mouse monoclonal anti-GFP at 1/1000 (clones 7.1 and 13.1, Roche), mouse monoclonal anti-TY at 1/250 (53), mouse anti-TgAtrx1 at 1/200 (39) and rabbit polyclonal anti-TgCpn60 at 1/2000 (43), mouse monoclonal anti-P21 at diluted 1/200 (T8 4G10) (79). Cysts were stained with a 1/300 dilution of a biotin-labelled *Dolichos biflorus* lectin (Sigma; L-6533) for 1h and revealed using a 1/300 dilution of FITC-conjugated streptavidin (Invitrogen; SNN1008). Staining of DNA was performed on fixed cells by incubating them for 5 min in a 1 μg/ml 4,6-diamidino-2-phenylindole (DAPI) solution. All images were acquired at the MRI facility from a Zeiss AXIO Imager Z2 epifluorescence microscope or a Zeiss lsm880 Fast Airyscan confocal microscope. Images were analyzed with by the ZEN software v2.5 (Zeiss) and ImageJ v1.53t (US National Institutes of Health). Z-stack acquisition and maximal intensity projection was systematically used to visualize the signal over the whole width of the cysts. Adjustments for brightness and contrast were applied uniformly on the entire image. Vacuoles or cysts were chosen randomly and identified based on the absence or presence of cyst wall labeling.

### Apicoplast loss quantification

The presence and the integrity of the apicoplast was evaluated through TgCpn60 staining and by comparing with the number of DAPI-stained nuclei in each cyst. When applicable (i.e. for small or medium-sized cysts) an ImageJ v1.53t (US National Institutes of Health) pipeline was used to quantify the signal for apicoplast to nucleus ratio calculation. Briefly, for both image stacks the DAPI and apicoplast signal channels, maximum intensity projection images were created and converted to 8 bits, and threshold was adjusted to create a binary mask version of the image, object counts were then generated through the ‘analyze particle’ function with the adequate size and circularity parameters.

### Immunoblot analysis

Protein extracts from freshly egressed tachyzoites or recently isolated bradyzoites were prepared in Laemmli sample buffer, separated by SDS-PAGE and transferred onto nitrocellulose membrane using the Mini-Transblot system (BioRad) according to the manufacturer’s instructions. Primary antibodies used in this study and their respective dilutions were: mouse monoclonal anti-GFP at 1/1000 (clones 7.1 and 13.1, Roche), mouse monoclonal anti-TY at 1/200 (53), mouse monoclonal anti-P21 at diluted 1/50 (T8 4G10) (79) and mouse anti-actin at 1/100 (80).

### Serological confirmation of infection by dot blot

Peripheral blood was collected intracardially without anticoagulant and centrifuged for 10 min at 14 000rpm. The serum was stored at -20°C. Tachyzoites were released from infected HFF, lysed in Laemlli buffer 1x (Biorad ref #1610737) + 5 % βmercapto-ethanol (BioRad ref #161-0710), and heated for 5 min at 70°C. Solubilized proteins were applied on a nitrocellulose membrane, which was incubated overnight at 4°C with each serum diluted at 1/1000. Immunologic detection was performed using horseradish peroxidase-conjuged anti-mouse IgG antibodies. Peroxidase activity was visualized by chemiluminescence and quantified using ChemiDoc camera (Biorad).

### Semiquantitative RT-PCR

Total RNA was extracted from *T. gondii* tachyzoites using the Nucleospin RNA II kit (Macherey-Nagel). Reverse transcription was performed with the Superscript III first-strand synthesis kit (Invitrogen). 1 µg of RNA was used as a template per RT reaction. Specific primers of *T. gondii glyceraldehyde 3-phosphate dehydrogenase 2* (*GAPDH2*, ML5049/ML5680) or *TgAPT1* (ML4097/ML4098) were used for subsequent PCR amplification with GoTaq polymerase (Promega) for twenty-five cycles as follows: 15s of denaturation at 95 °C, 30s of annealing at 54 °C and 40s of elongation at 68°c. Similar volumes of samples for which no reverse transcriptase was included in the initial cDNA preparation were used as a control for genomic DNA contamination.

### Statistical analyses

Statistical analyses (unpaired Student’s t-test for comparisons between two groups, one-way ANOVA with post-hoc Tukey test for comparisons between three or more groups, Mann-Whitney non-parametric test for cyst size comparison, and Mantel–Cox logrank test for survival assays) were performed with Prism v8.3 (Graphpad). Unless specified, values are expressed as means + standard deviation (SD).

## Acknowledgments

We are grateful to B. Striepen, D. Soldati-Favre and M. Parsons for providing plasmids and antibodies and H. Acha-Orbea for the DC cell line. We also thank M. Gissot and J.-F. Dubremetz for advice with the cyst purification protocol. Thanks to the MRI imaging and cytometry facilities for granting access to their equipment and thanks to C. Duperray and Y. Bordat for help with formatting of the flow cytometry results. We thank R. Balouzat, R. Ecalard and the zootechnicians at Inserm UMS006-CREFRE mouse facility, and F. L’Faqihi-Olive, V. Duplan-Eche, A.-L. Iscache, and H. Garnier for assistance at the Infinity flow cytometry facility. We also thank the developers and the managers of the VeupathDB.org/ToxoDB.org databases, as well as scientists who contributed datasets.

This work was supported by a grant from the Agence Nationale de la Recherche (ANR-19-CE15-0023) to N.B. and S.B. and the PIA PARAFRAP Consortium (ANR-11-LABX0024) to N.B.

