## Supplemental Figs 1-10, Supplemental Table 1 for "The apicoplast is important for the viability and persistence of *Toxoplasma gondii* bradyzoites"

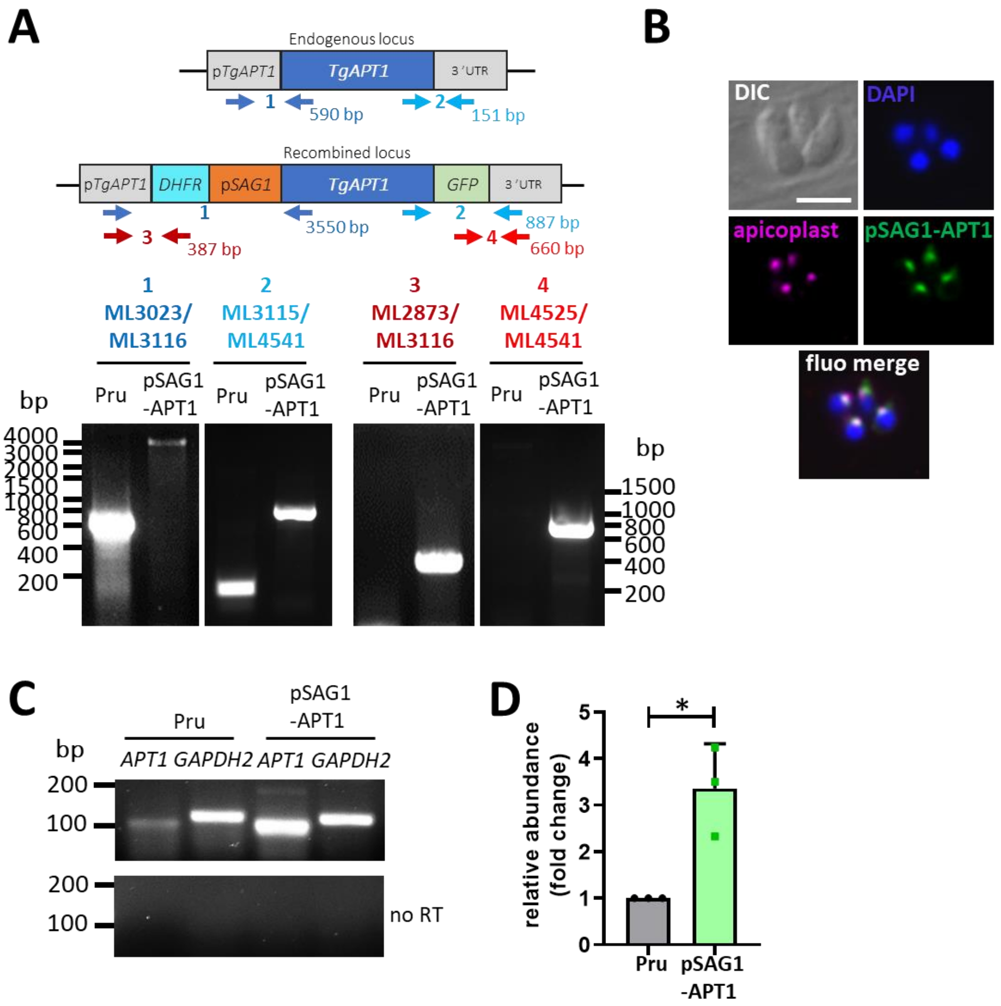


**Figure S1. Using a *SAG1* promoter to express the GFP-fused APT1 protein does not affect protein localization but leads to increased mRNA expression at the tachyzoite stage.** A) Diagnostic PCRs show proper replacement of the native *APT1* gene by the *SAG1* promoter-driven APT1-GFP fusion cassette. Primer positions and expected size of the fragments are mentioned on the schematic representations of the wild-type and recombined loci. B) Immunofluorescence assays shows that GFP-fused APT1 localizes to the apicoplast (labelled here with membrane marker Atrx1). DNA was labelled with DAPI. Scale bar=5 µm C) Semi-quantitative RT-PCR analysis performed with primers specific of *APT1* or of the internal control *glyceraldehyde 3-phosphate dehydrogenase 2* (*GAPDH 2*). Control reactions where the reverse transcriptase was omitted are also shown ('no RT'). D) Quantification of the RT-PCRs (performed as described in C) was done by band densitometry and normalized with the internal control. Shown are means +SD of *n*=3 independent experiments. * $p \leq 0.05$, unpaired Student's t test.

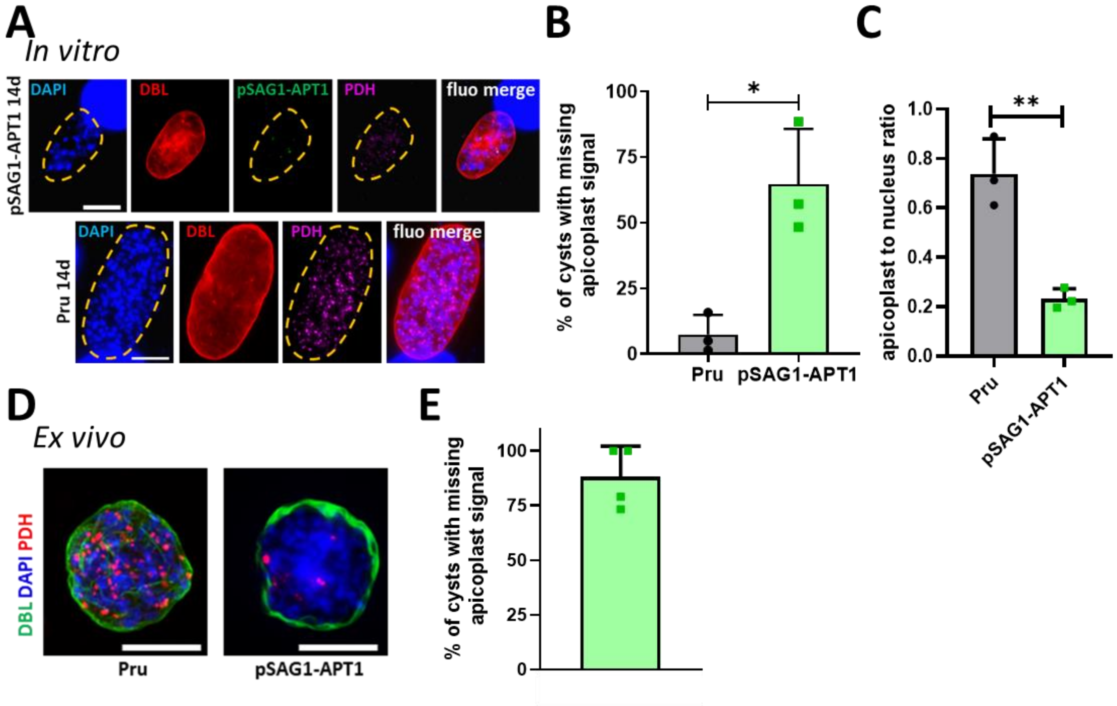


**Figure S2. APT1 depletion also impacts the PDH-E2 apicoplast marker**. A) *In vitro*-generated cysts (14 days of alkaline pH differentiation) leads to the disappearance of both APT1 and of the PDH-E2 apicoplast marker (magenta) in mature cysts labelled with DBL (red). Scale bar=10 µm. B) The integrity of the PDH-E2-labelled apicoplast signal was quantified after 14 days of alkaline pH-induced differentiation and showed a significant loss in the pSAG1-APT1 cell line. Data are mean from *n*=3 independent experiments +SD. * $p \leq 0.05$, unpaired Student's t-test. C) When applicable, fluorescent signal for individual nuclei (DAPI-stained) and PDH-E2-stained apicoplasts was recorded within cysts, quantified, analyzed and plotted to give an apicoplast to nucleus value. Shown are mean +SD from *n*=3 independent experiments. ** $p \leq 0.01$, unpaired Student's t-test. D) Labelling of mouse brain-derived cysts with anti-PDH-E2 shows partial loss of the signal in the pSAG1-APT1 cell line. Scale bar= 10 µm. E) Quantification of PDH-E2 signal in brain-derived pSAG1-APT1 cysts shows that most display at least a partial apicoplast loss 40 days post-infection. Shown are the mean +SD from *n*=4 different mice from a single *in vivo* experiment; 13 to 19 cysts were counted in each dataset.

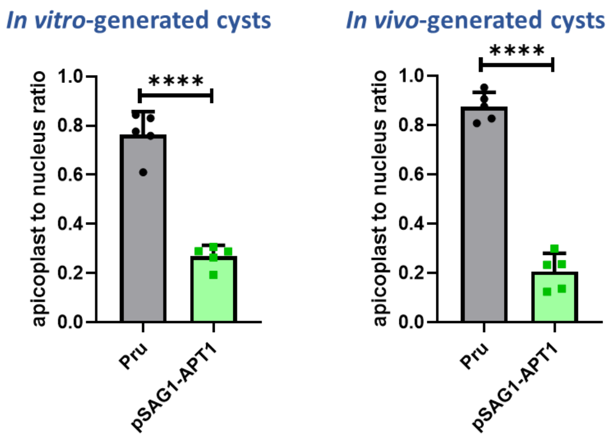


**Figure S3. APT1 depletion leads to a considerable loss of apicoplast signal within cysts.** When applicable, fluorescent signal for individual nuclei (DAPI-stained) and apicoplasts (Cpn60-labelled) was recorded within cysts, quantified, analyzed and plotted to give an apicoplast to nucleus value. Data for *in vitro*-generated cysts were acquired from *n*=5 independent series including 22 to 56 cysts; Data for *ex vivo*-generated cysts were acquired from *n*=5 independent series including 7 to 16 cysts. Represented are the means +SD. **** $p \leq 0.0001$, unpaired Student's t-test.

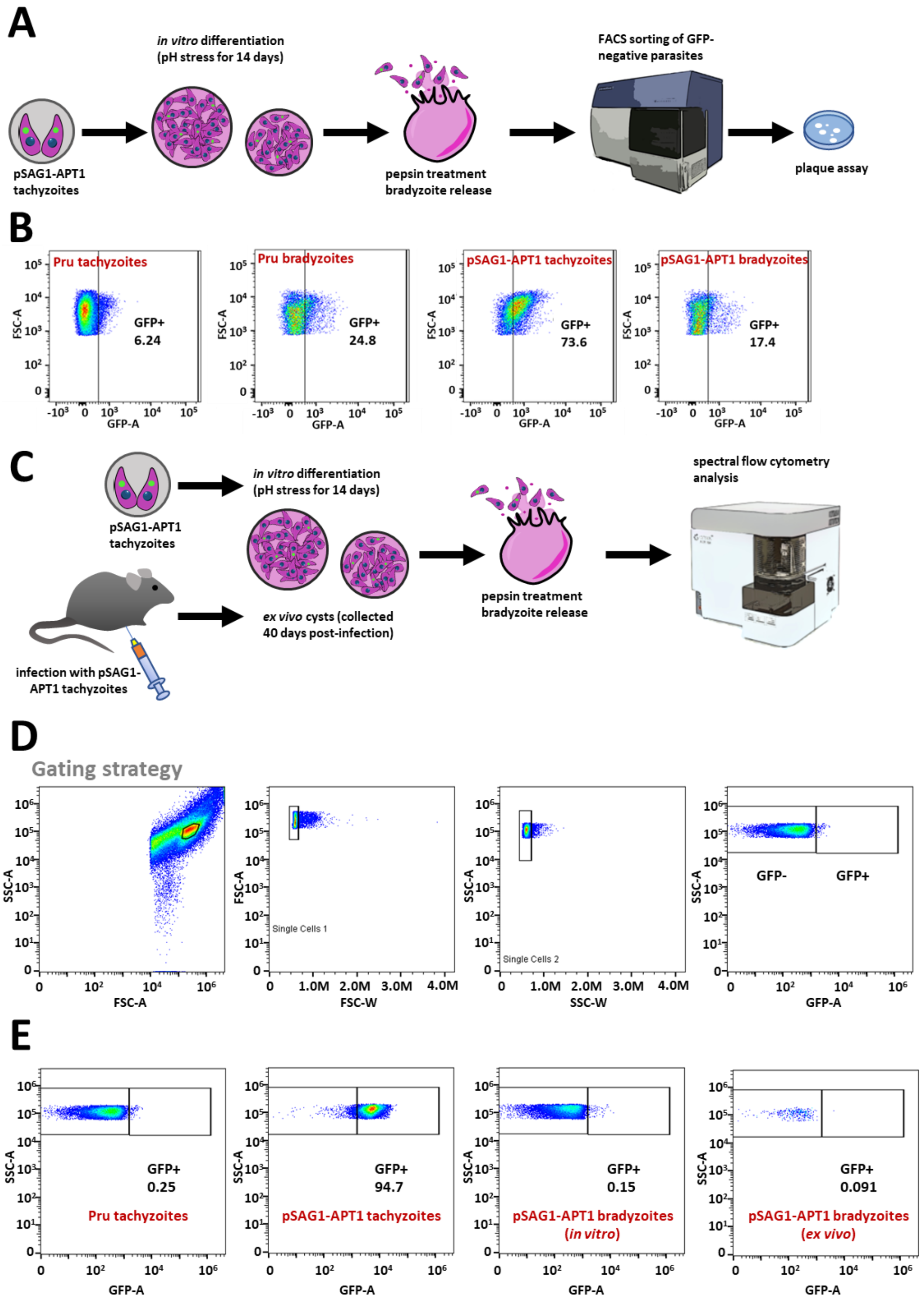


**Figure S4. Isolation and cytometric analysis of pSAG1-APT1 bradyzoites.** A) Schematic representation of the strategy for isolating *in vitro*-differentiated APT1-deficient pSAG1-APT1 bradyzoites, which were sorted on the basis of an absence of GFP fluorescence and plated on host cells monolayers for plaque assay. B) Typical flow cytometry profiles of tachyzoites and *in vitro*-

differentiated bradyzoites of the pSAG1-APT1 and Pru control cell lines. There is a marked decrease in GFP fluorescence in the pSAG1-APT1 bradyzoites compared with tachyzoites. GFP-negative Pru and pSAG1-APT1 bradyzoites were used for subsequent plaque assays. C) *In vitro*- and *ex vivo*-derived bradyzoites were also analyzed by spectral flow cytometry. D) Gating strategy for isolating singlet parasites for GFP signal measurement. E) GFP fluorescence profiles of tachyzoites or *in vivo*- and *ex vivo*-derived bradyzoites.

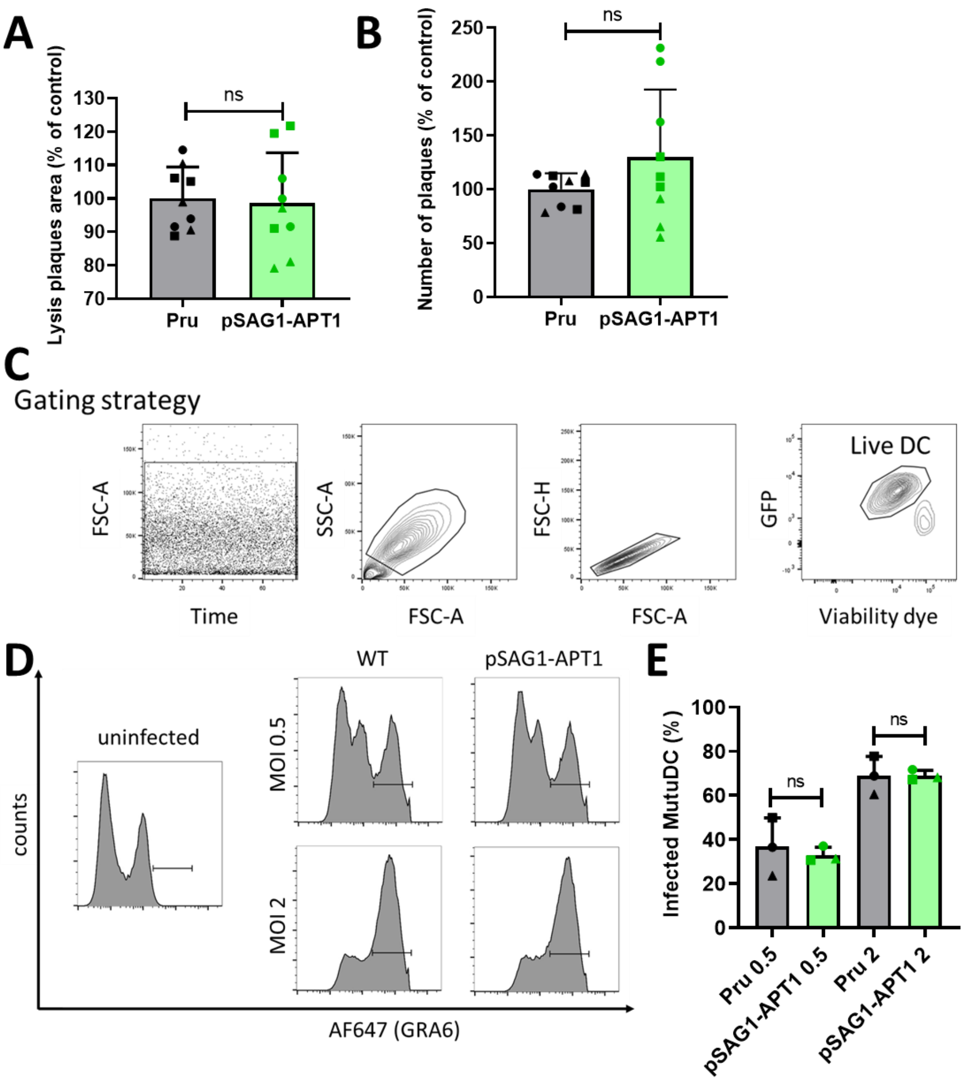


**Figure S5. pSAG1-APT1 tachyzoites have no fitness defect or advantage in *in vitro* models of infection.** Quantification of lysis plaque area (A) or number (B) after infection of HFF host cells by the pSAG1-APT1 and Pru cell lines. Values are means of nine values (representing *n*=3 independent experiments, containing three technical triplicates) +SD. Symbols are matched between identical experimental groups. Mean value for the Pru cell line was set to 100% as a reference. 'ns' not significant, unpaired Student's t test. C) Gating strategy for the dendritic cell (DC) infection assay. To measure the rate of infected DC, successive gates were placed to analyze singlet live GFP+ MutuDC. The proportion of AF647+ DC, corresponding to cells in which GRA6 is secreted into the parasitophorous vacuole and thus detected by intracellular staining, was calculated at a multiplicity of infection (MOI) of 0.5 and 2. The gate was defined using uninfected cells as a negative control (D). E) Quantification of DC infection by the pSAG1-APT1 and Pru cell lines. Mean values +SD representing quantification from *n*=3 independent experiments are shown. 'ns' not significant, unpaired Student's t test.

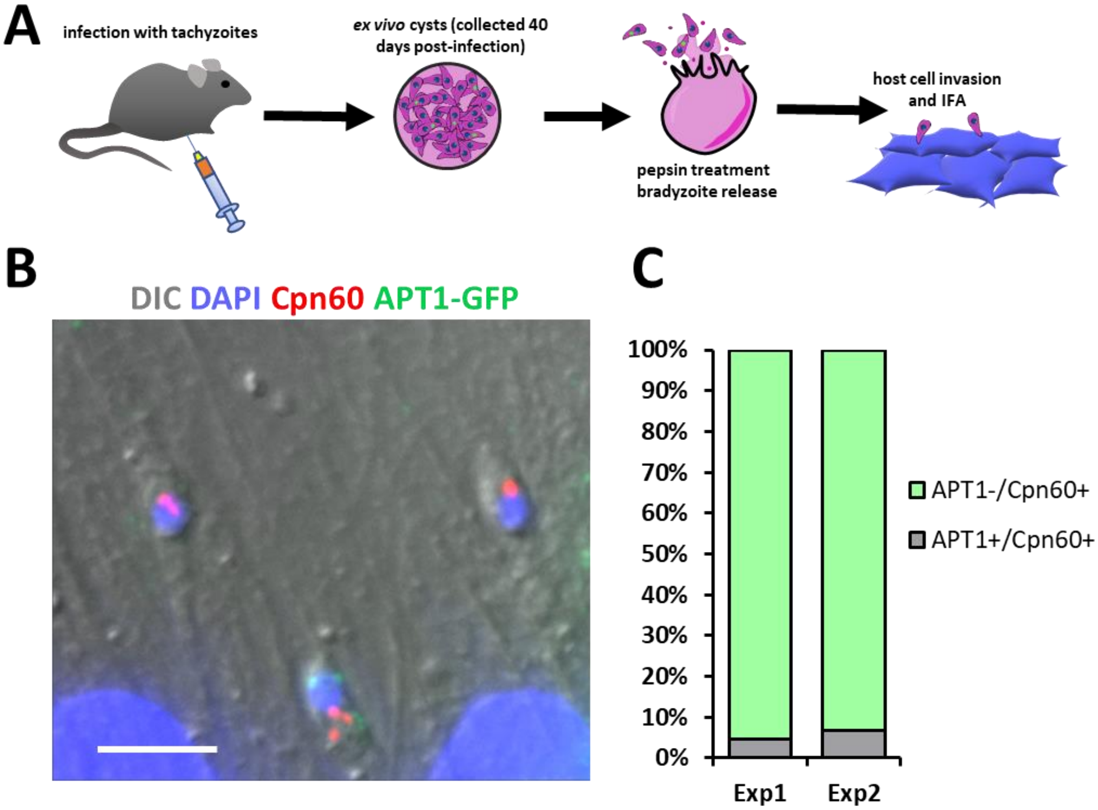


**Figure S6. Viable *in vivo*-derived pSAG1-APT1 bradyzoites still harbor an apicoplast.** A) pSAG1-APT1 parasites were used to infect mice and 40 days post-infection cysts were recovered and bradyzoites were extracted to be used subsequently to invade host cells for 3 hours and then further processed for immunofluorescence assay (IFA) in order to detect the presence of the apicoplast. B) Typical merged image of intracellular bradyzoites fixed 3 hours post-invasion and showing the presence of the apicoplast marker Cpn60 (red) in spite of the absence of APT1-GFP (green). DNA was labelled with DAPI (blue). DIC: differential interference contrast. Scale bar=5 µm. C) Quantification of the presence of the APT1-GFP and Cpn60 (apicoplast) signals in freshly invaded *in vivo*-generated bradyzoites. Represented data are from two independent experiments with counts from 43 (Exp.1) and 30 (Exp. 2) individual parasites.

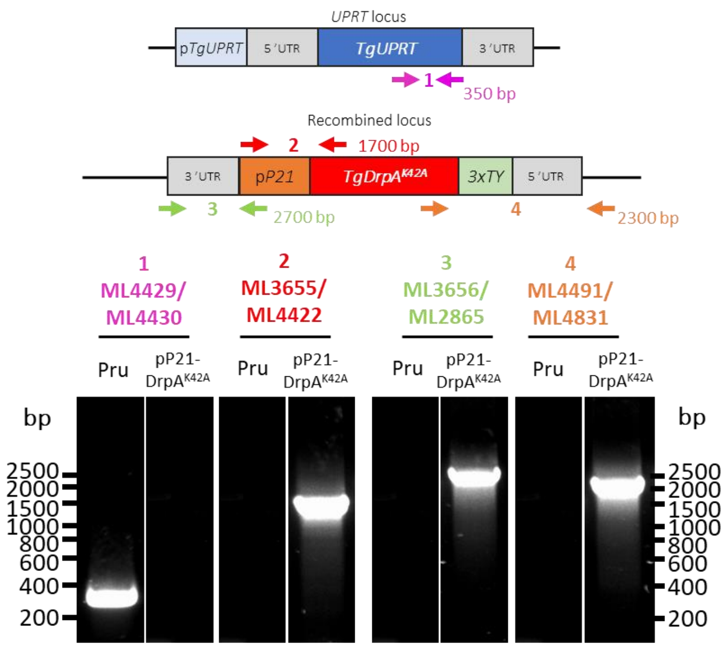


**Figure S7. Insertion of a mutated copy of the *DrpA* gene at the *UPRT* locus.** Diagnostic PCRs show proper replacement of the native *UPRT* gene by the *P21* promoter-driven DrpA$^{K42A}$-triple TY tag fusion cassette. Primer positions and expected size of the fragments are mentioned on the schematic representations of the wild-type and recombined loci.

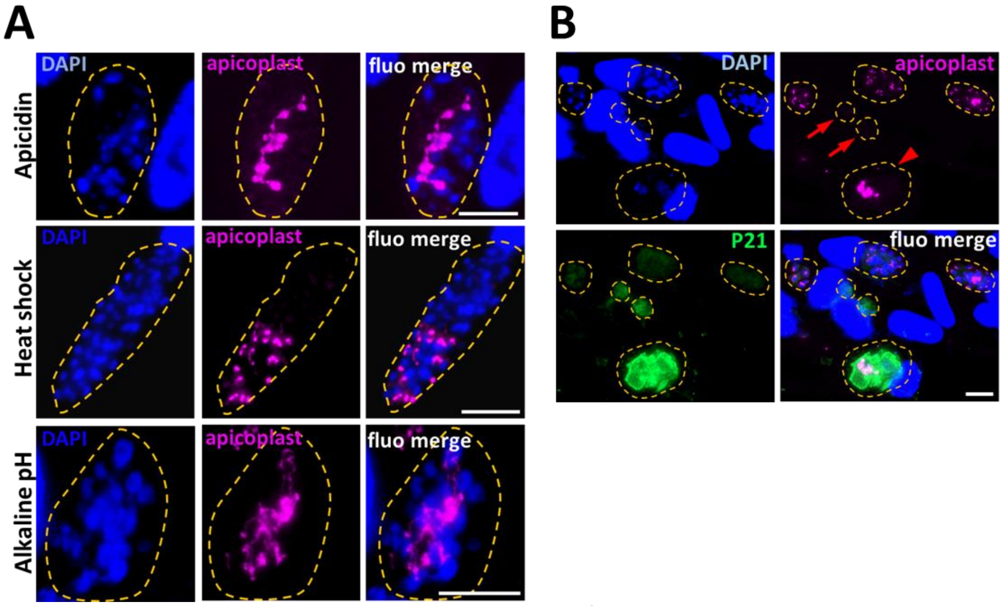


**Figure S8.** ***In vitro*-differentiated pP21-DrpA$^{K42A}$ bradyzoites show defects in plastid segregation.** A) Immunofluorescence analysis of DBL-labelled cysts obtained after six days of apicidin treatment, heat shock or three days of alkaline pH stress. The apicoplast was labelled with the Cpn60 marker and shows a bulky/elongated pattern in some parasites and is absent from others. Cyst wall is outlined with dashed lines. DNA was stained with DAPI. Scale bar = 10 µm. B) Apicidin-treated parasites displaying the most severe apicoplast morphological defects, like organelle reticulation and segregation problems (arrowhead) or loss of the organelle (arrows), are those with a higher P21 expression. Vacuoles/cyst walls are outlined with dashed lines. DNA was stained with DAPI. Scale bar = 10 µm.

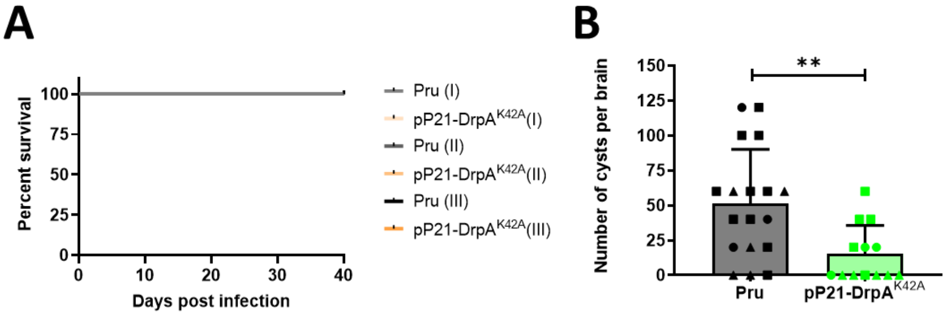


**Figure S9. Mouse infection with the pP21-DrpA$^{K42A}$ parasite line shows that it is does not induce a strong acute phase and generates a low cyst burden.** A) Survival curves of CBA mice infected with 100 (experiment I) or 2000 parasites (experiment II and III) of the Pru or pP21-DrpA$^{K42A}$ cell lines shows that the latter does not induce mortality during the acute phase. Data from *n*=3 independent experiments are plotted. B) Quantification of pP21-DrpA$^{K42A}$ cyst burden in infected mice. Cyst numbers were estimated based on microscopic counts. Values are from *n*=3 independent experiments and are expressed as means +SD. Symbols are matched between identical experimental groups. ** $p \leq 0.01$, unpaired Student's t test.

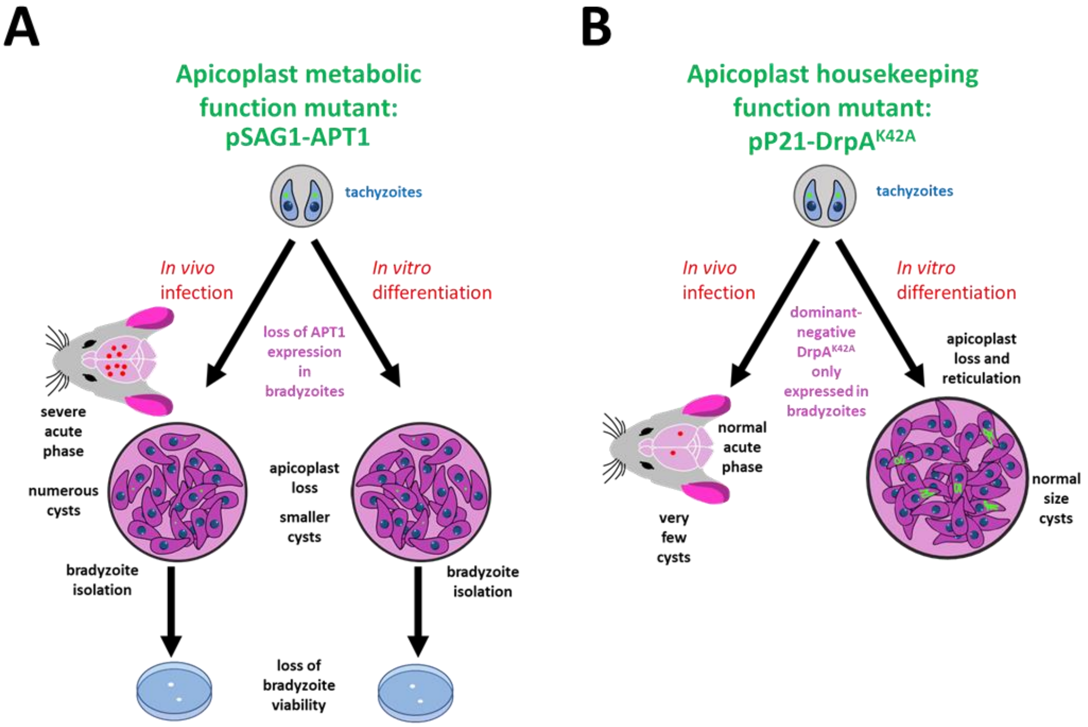


**Figure S10. Summary of the phenotypes associated with the two apicoplast conditional mutants generated in this study.** Schematic representation of the phenotypic outcomes that shows for instance that while APT1-deficient parasites generate numerous but small cysts containing essentially non-viable bradyzoites, and parasites for which DrpA function is affected have a strong defect in cystogenesis. Yet, both highlight the importance of the apicoplast for the persistence and the development of the parasites in the host.

**Table S1.** Oligonucleotides used in this study.

| **Primer name** | **Primer sequence** |
|---|---|
| ML2087 | AAGTGCTCCCACGTCCCTCACCAT |
| ML2088 | AAAAATGGTGAGGGACGTGGGAGC |
| ML2865 | CTCGTCTCCGGTGGAATGA |
| ML2873 | GGATCTGCACACCTGGTCTCGATG |
| ML3012 | TACTTCCAATCCAATTTAATGCTTAATTAAATGGAGGAATCGAAACGCTT |
| ML3013 | TCCTCCACTTCCAATTTTAGCTCCGTACTTGGTCTTCGAGAGAG |
| ML3015 | AAATTAATTAAGCCTTTGGCTCCTGAGA |
| ML3016 | AAATTAATTAAACAACCGTGTGTTTACACGA |
| ML3023 | CTCGCCTGCGACTCTAAATCC |
| ML3026 | AAGTTGACGGATAATAAACATGACGAG |
| ML3027 | AAAACTCGTCATGTTTATTATCCGTCA |
| ML3030 | AAGTTGAATACATGCGTCAACTTGGGG |
| ML3031 | AAAACCCCAAGTTGACGCATGTATTCA |
| ML3088 | CACAGGGTGGCGACTGGTTGAGGAATCACAATGAAGATCCGATCTTGCTG |
| ML3089 | AACGAGGTCTCTTCTCCTTCGTCATGTTTATTACTTGTACAGCTCGTCCA |
| ML3115 | TCGCGTCTGTCCTCTTCTTCC |
| ML3116 | GTCTACGACCTCTTTTCCGTCTCA |
| ML3445 | AAGTCAGGGCTTCTAAAATGGCGC |
| ML3446 | AAAAGCGCCATTTTAGAAGCCCTG |
| ML3655 | TCAGCGGCCGCCTCTCCTTGTCTTCTTACCG |
| ML3656 | ATAAGATCTTTTGAAGCTAGGCTGCG |
| ML4097 | GCGGTGATTTCAATGGGCTG |
| ML4098 | GACCTGGTGAAGAGCCAGTC |
| ML4417 | TTTTGATCAATGGAGGAGTTGATTCCTG |
| ML4418 | TTTCCTAGGTCGACAAGCGGAGGCG |
| ML4419 | CTAGGGAAGTTCATACTAATCAAGATCCACTAGATCTAGAAGTTCATACTAATCAAGATCCACTA[illegible] |
| ML4420 | CTAGTATCTAGTGGATCTTGATTAGTATGAACTTCTAGATCTAGTGGATCTTGATTAGTATGAAC[illegible] |
| ML4421 | CAGAGTGTGGGGGCGTCTTCTGTGCTC |
| ML4422 | GAGCACAGAAGACGCCCCCACACTCTG |
| ML4429 | ACTGCGACGACATACTGGAGAAC |
| ML4430 | AAGAAAACAAAGCGGAACAACAA |
| ML4491 | GAGCTCGTTGCTCAGCTGTATCGAGAAG |
| ML4525 | TGGACGGCGACGTAAACGG |
| ML4541 | GGTCTCTTCTCCTTCGTCATG |
| ML4831 | TAGAACTCGCACACCCCTCG |
| ML5049 | AAAAGATTGCGGGACGCAGT |
| ML5680 | CGTTCCATCGCAGCACGG |
| TOX11 | GCGTCGTCTCGTCTAGATCG |
| TOX9 | AGGAGAGATATCAGGACTGTAG |
